# Epigenetic reprogramming of plasmacytoid dendritic cells drives type I interferon-dependent differentiation of acute myeloid leukemias for therapeutic benefit

**DOI:** 10.1101/2020.08.23.235499

**Authors:** Jessica M. Salmon, Izabela Todorovski, Stephin J. Vervoort, Kym L. Stanley, Conor J. Kearney, Luciano Martelotto, Fernando Rossello, Tim Semple, Gisela Mir-Arnau, Magnus Zethoven, Michael Bots, Eva Vidacs, Kate McArthur, Elise Gressier, Nicky de Weerd, Jens Lichte, Madison J. Kelly, Leonie Cluse, Simon J. Hogg, Paul J. Hertzog, Lev Kats, Daniel D. de Carvalho, Stefanie Scheu, Sammy Bedoui, Benjamin T. Kile, Andrew Wei, Pilar M. Dominguez, Ricky W. Johnstone

## Abstract

Pharmacological inhibition of epigenetic enzymes can have therapeutic benefit, particularly against hematological malignancies. While these agents can affect tumor cell growth and proliferation, recent studies have demonstrated that pharmacological de-regulation of epigenetic modifiers may additionally mediate anti-tumor immune responses. Here we discovered a novel mechanism of immune regulation through the inhibition of histone deacetylases (HDACs). In a genetically engineered model of t(8;21) AML, leukemia cell differentiation and therapeutic benefit mediated by the HDAC inhibitor panobinostat required activation of the type I interferon (IFN) signaling pathway. Plasmacytoid dendritic cells (pDCs) were identified as the cells producing type I IFN in response to panobinostat, through transcriptional activation of IFN genes concomitant with increased H3K27 acetylation at these loci. Depletion of pDCs abrogated panobinostat-mediated activation of type I IFN signaling in leukemia cells and impaired therapeutic efficacy, while combined treatment of panobinostat and recombinant IFNα improved therapeutic outcomes. These discoveries offer a new therapeutic approach for t(8;21) AML and demonstrate that epigenetic rewiring of pDCs enhances anti-tumor immunity, opening the possibility of exploiting this cell type as a new target for immunotherapy.

## INTRODUCTION

Two of the most prevalent cytogenetic subtypes of primary AML, t(8;21) and inv(16), involve genes encoding core binding factors (CBFs), AML1 and CBFbeta, respectively (1). Although CBF-associated AML is considered “favourable risk”, close to 50% of patients will relapse at 5 years and die of their disease (2). The t(8;21) translocation produces the AML1-ETO fusion oncoprotein, which affects the normal transcriptional activity of AML1 (3,4). In addition, mutations in N/KRAS frequently co-occur in t(8;21) AML patients (5). We previously established a mouse model with pathological and molecular features of human AML, initiated by the constitutive co-expression of AML1-ETO9a (containing a transcript with alternative splicing in exon 9 of *ETO*) and an oncogenic allele of NRAS (NRAS^G12D^) in mouse hemopoietic progenitor cells (6,7). We and others previously showed that AML1-ETO can recruit histone deacetylases (HDACs) (7-9), resulting in repression of AML1-target genes and suppression of hemopoietic cell differentiation to promote leukemogenesis. Importantly, treatment with HDAC inhibitors (HDACis) induced terminal differentiation of leukemia cells and enhanced survival in t(8;21)AML mouse models (7).

While a number of HDACis have been FDA approved for hematological malignancies (10), their therapeutic impact as single agents has been modest. Studies using genetically and histologically distinct pre-clinical cancer models demonstrated that the efficacy of HDACis was reduced in immune compromised recipient mice or following immune cell depletion in wild-type mice, indicating an immune dependent component in their mechanism of action (11). In line with this, recent studies have shown promising results of combining HDACi with immunotherapy (12). While these reports point to the ability of HDACi to engage the host immune system, exactly how this is achieved remains ill-defined. Previous studies have focused on the ability of HDACis to enhance the immunogenicity of tumor target cells by promoting the expression of key immune-modulating genes that are epigenetically silenced during tumor immune evasion (13,14). However, the impact of these agents on the activity of other cells in the tumor microenvironment, in particular cells that regulate host and/or tumor immunity, has been less well studied.

Here we investigated the genes and molecular pathways altered by HDACis in a model of t(8;21) AML. Our results reveal that the HDACi panobinostat induced plasmacytoid dendritic cells (pDCs) to increase expression of type I interferon (IFN). The combined effect of panobinostat and pDC-derived IFN resulted in leukemia cell differentiation *in vivo* and a pronounced therapeutic outcome. Perturbation of this combined effect through genetic deletion of the type I IFN receptor (IFNAR) on the leukemia cells or depletion of pDCs *in vivo* significantly abrogated the therapeutic effects of panobinostat. Moreover, the combinatorial effects of panobinostat and type I IFN could be therapeutically exploited through the addition of recombinant IFNα to the panobinostat therapy regimen that resulted in greatly enhanced therapeutic benefit. This is the first study demonstrating that HDACis can mediate profound anti-tumor responses through immune modulatory effects on pDCs resulting in enhanced type I IFN gene transcription. The resultant type I IFN is necessary for HDACi-induced leukemia differentiation and therapeutic benefit, identifying a unique molecular interplay between effects of epigenetic agents on the tumor target cells and host immune cells.

## RESULTS

### Panobinostat induces differentiation of AML cells through activation of the type I interferon pathway

A pre-clinical mouse model of t(8;21) AML developed through retroviral expression of AML1-ETO9a and NRAS^G12D^ (A/E9a;NRAS^G12D^) in hematopoietic cells (6,7) was utilized to study the therapeutic effects of HDACis and decipher the molecular events that underpinned their anti-cancer activities (**Suppl Fig. 1A**). Treatment of mice bearing A/E9a;NRAS^G12D^ AML with panobinostat resulted in reduced tumor burden compared to vehicle-treated animals as assessed by reduced GFP^+^ cells in the peripheral blood (**Fig. 1A**) and a decrease in bioluminescence (**Fig. 1B**). The panobinostat-induced reduction in leukemia cell numbers manifested in a significant median survival advantage of 38.5 days (**Fig. 1C**). We previously showed that treatment of A/E9a;NRAS^G12D^ leukemias with panobinostat resulted in leukemia cell cycle arrest and myeloid differentiation (7,15). To determine the minimum panobinostat treatment time required for t(8;21) AML cells to undergo differentiation, leukemia cells were harvested from the bone marrow over the first week of therapy for assessment. BrdU incorporation assays demonstrated that panobinostat reduced leukemia cell proliferation from day 3 onwards with only approximately 10% of cells in the panobinostat cohort in S-phase after 5 days of therapy compared to 40% of cells in S-phase in the vehicle-treated group (**Fig. 1D**). Consistent with our previous studies (6,7), panobinostat treatment resulted in decreased expression of c-Kit and increased expression of CD11b (**Fig. 1E**), indicating terminal myeloid differentiation towards the granulocyte lineage.

**FIGURE 1:**
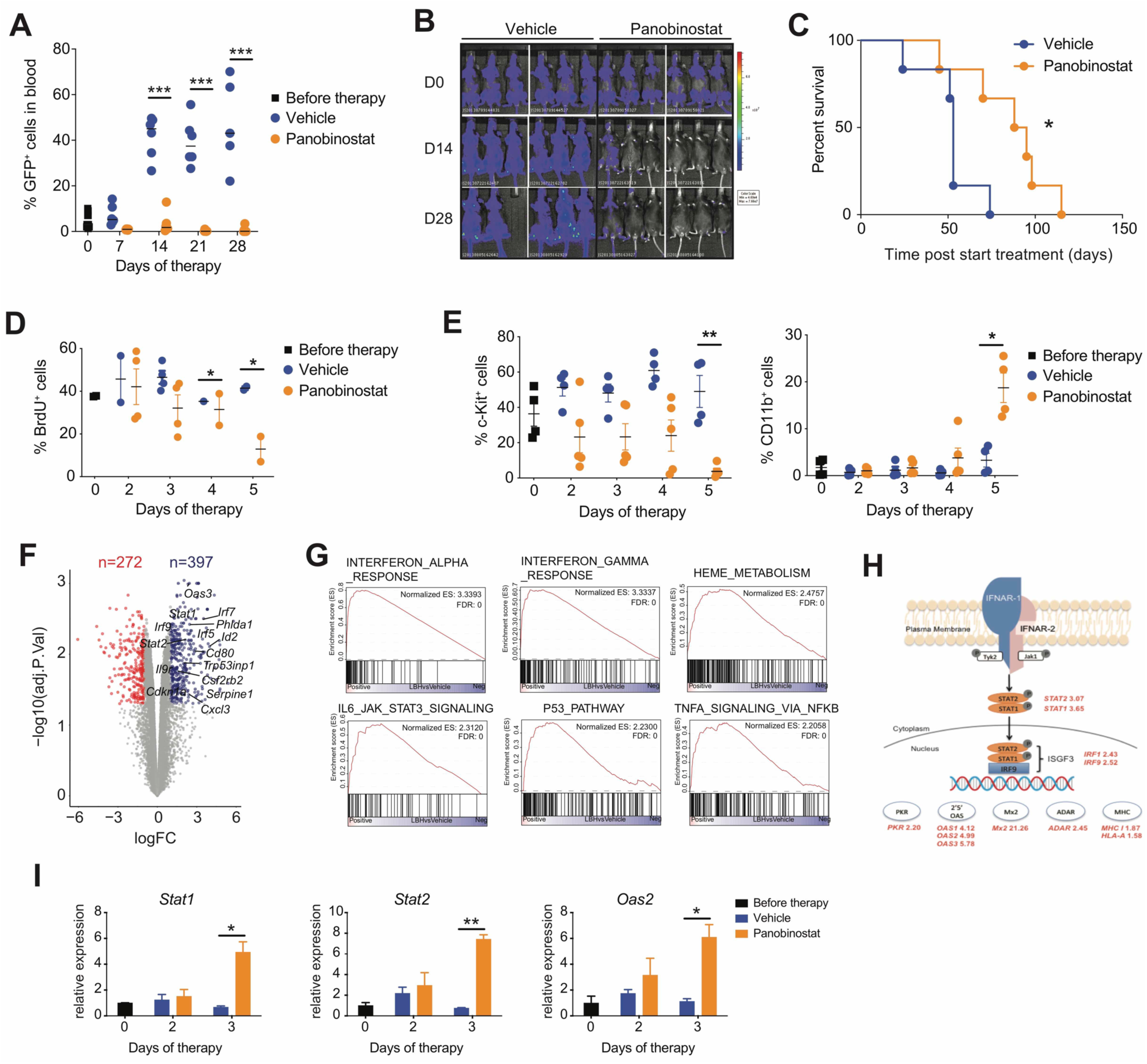
Panobinostat induces differentiation of AML cells through activation of the type I interferon pathway. (**A**) Percentage of GFP positive cells in the peripheral blood of mice bearing A/E9a;NRAS^G12D^-driven leukemias treated with vehicle or panobinostat (n=5-6 mice/group; \*\*\**P*<0.001). (**B**) Bioluminescence imaging of individual tumor-bearing animals over the course of therapy with vehicle or panobinostat. (**C**) Kaplan-Meier survival curves of mice bearing A/E9a;NRAS^G12D^-driven leukemias treated with vehicle or panobinostat (n=6 mice/group; \**P*<0.05). (**D**) Cell cycle analysis of A/E9a;NRAS^G12D^ leukemia cells within the bone marrow of animals treated with vehicle or panobinostat. The percentage of cells in S-phase (BrdU-positive) was determined by flow cytometry (n=3; \**P*<0.05). (**E**) Flow cytometric analysis of the cell surface expression of c-Kit and CD11b on tumor cells in the bone marrow of A/E9a;NRAS^G12D^ tumor-bearing mice treated for 5 days with vehicle or panobinostat (n=3; \**P*<0.05, \*\**P*<0.005). (**F**) Volcano plot of the DEG (logFC>1; adj.Pval<0.05) between A/E9a;NRAS^G12D^ cells isolated from panobinostat-treated and vehicle-treated leukemia-bearing mice. (**G**) GSEA of upregulated genes shown in (F). (**H**) IFNAR signaling pathway and increased fold change in individual genes induced by panobinostat treatment. (**I**) Expression by qRT-PCR of *Stat1, Stat2* and *2’5’Oas* on tumor cells isolated from vehicle- and panobinostat-treated animals over 3 days of therapy (n=3; \**P*<0.05, \*\**P*<0.005).

To determine the transcriptional changes that might underpin the induction of differentiation of leukemia cells following exposure to panobinostat, RNA sequencing (RNA-seq) was performed on leukemias harvested from tumor-bearing mice treated with either vehicle or panobinostat for 3 days, the point at which panobinostat-treated AML cells are demonstrating features of differentiation. Principal component analysis showed a clear separation between the transcriptional profiles of leukemia cells from panobinostat-treated or vehicle-treated (control) mice (**Suppl Fig. 1B**). A supervised analysis identified 669 differentially expressed genes (DEG) in leukemias harvested from panobinostat-treated and control mice (log fold change (logFC)> 1, adj.Pval<0.05), the majority of which were upregulated after panobinostat treatment (n=397, **Fig. 1F** and **Suppl Table 1**). Pathway analysis of these upregulated genes through gene set enrichment analysis (GSEA) and gene ontology (GO) enrichment revealed significant enrichment of genes associated with type I IFN and anti-viral responses, as well as IFN gamma (IFNγ) response, p53 pathway and cytokine signaling, including IL-6 and TNFα (**Fig. 1F and G and Suppl Fig. 1C**). Upregulation of transcription factors primarily involved in type I IFN signaling (*Stat1, Stat2, Irf1, Irf9*) was prevalent and a concomitant increase in expression of STAT target genes *Mx2* and *2’5’Oas* was observed (**Fig. 1F, H** and **Suppl Table 1**). qRT-PCR was employed as an orthogonal method to confirm the increased expression of *Stat1, Stat2* and *Oas2* in leukemia cells exposed to panobinostat for 3 days *in vivo* (**Fig. 1I**).

### Expression of IFNAR1 on A/E9a;NRAS^G12D^ leukemia cells is critical for the efficacy of panobinostat *in vivo*

Type I IFN signaling through IFNAR expressed on tumor cells can induce cell cycle arrest and cell death (16,17). To determine the functional and therapeutic relevance of type I IFN signal activation following exposure of A/E9a;NRAS^G12D^ AML to panobinostat, leukemias with genetic knockout of *IFNAR1* were developed by transducing fetal liver cells harvested from *Ifnar1*^*-*/-^ mice (18) with retroviral constructs expressing AML1-ETO9a and NRAS^G12D^ (**Fig. 2A**). Strikingly, when wild typemicetransplantedwith A/E9a;NRAS^G12D^;*Ifnar1*^-/-^ leukemias were treated with panobinostat, the therapeutic benefit of panobinostat observed in mice bearing A/E9a;NRAS^G12D^ AML was lost (**Fig. 2B**). This result was further confirmed in two additionalindependentlyderived A/E9a;NRAS^G12D^;*Ifnar1*^-/-^ leukemias (**Suppl Fig. 2A**). A readout for panobinostat activity in A/E9a;NRAS^G12D^leukemias is degradation of the AML1-ETO9a fusion protein (6,7) and this effect was also observed in panobinostat-treated A/E9a;NRAS^G12D^;*Ifnar1*^-/-^ cells, indicating that certain biochemical effects mediated by panobinostat remained unaffected by knockout of *Ifnar1* (**Fig. 2C**). Moreover, panobinostat caused a decrease in BrdU incorporation in both A/E9a;NRAS^G12D^and A/E9a;NRAS^G12D^;*Ifnar1*^-/-^ leukemias following treatment of tumor-bearing mice with panobinostat for 3 days (**Fig. 2D**). The ability of A/E9a;NRAS^G12D^;*Ifnar1*^-/-^ tumors to differentiate in response to panobinostat was determined through assessment of cell surface expression of c-Kit on tumor cells within the bone marrow over 4 days of therapy. There was no alteration in c-Kit expression on panobinostat-exposed A/E9a;NRAS^G12D^;*Ifnar1*^-/-^ cells in contrast to the significant decrease in c-Kit expression observed in A/E9a;NRAS^G12D^ leukemias exposed to panobinostat (**Fig. 2E**). Moreover,exposure of A/E9a;NRAS^G12D^;*Ifnar1*^-/-^ leukemias to panobinostat *in vivo* did not induce morphological changes consistent of myeloid differentiation such as condensation of nuclei and formation of cytoplasmic granules seen in panobinostat-treated A/E9a;NRAS^G12D^ cells (**Fig.2F**).RNA-seq on A/E9a;NRAS^G12D^;*Ifnar1*^-/-^ leukemias harvested from mice treated with panobinostat for 3 days was performed to determine if IFNAR deficiency affected the type I IFN response following treatment of A/E9a;NRAS^G12D^ cells with panobinostat (**Suppl Fig. 2B**). This revealed 534 DEG between panobinostat-treatedandvehicle-treated A/E9a;NRAS^G12D^;*Ifnar1*^-/-^ tumor cells (logFC>1, adj.Pval<0.05; **Fig. 2G and Suppl Table 2**). Comparison of the transcriptional changes observed in panobinostat-treated A/E9a;NRAS^G12D^and A/E9a;NRAS^G12D^;*Ifnar1*^-/-^ leukemias revealed that transcripts related to type I IFN signaling pathway, such as *Irf7, Irf9, Stat1* and *Stat2*, were only upregulated in A/E9a;NRAS^G12D^ cells (**Fig. 2H**). GSEA and GO analysis of the 361 upregulated genes in panobinostat-treated A/E9a;NRAS^G12D^;*Ifnar1*^-/-^ cells showed enrichment in p53 pathway, IFNγ response and cytokine signaling (**Fig. 2G and I and Suppl Fig 2C**), similar to A/E9a;NRAS^G12D^ cells (**Fig. 1G**). However pathways induced by type I IFN, observed in the panobinostat treated A/E9a;NRAS^G12D^ leukemias, were not found in treated A/E9a;NRAS^G12D^;*Ifnar1*^-/-^ cells.

**FIGURE 2:**
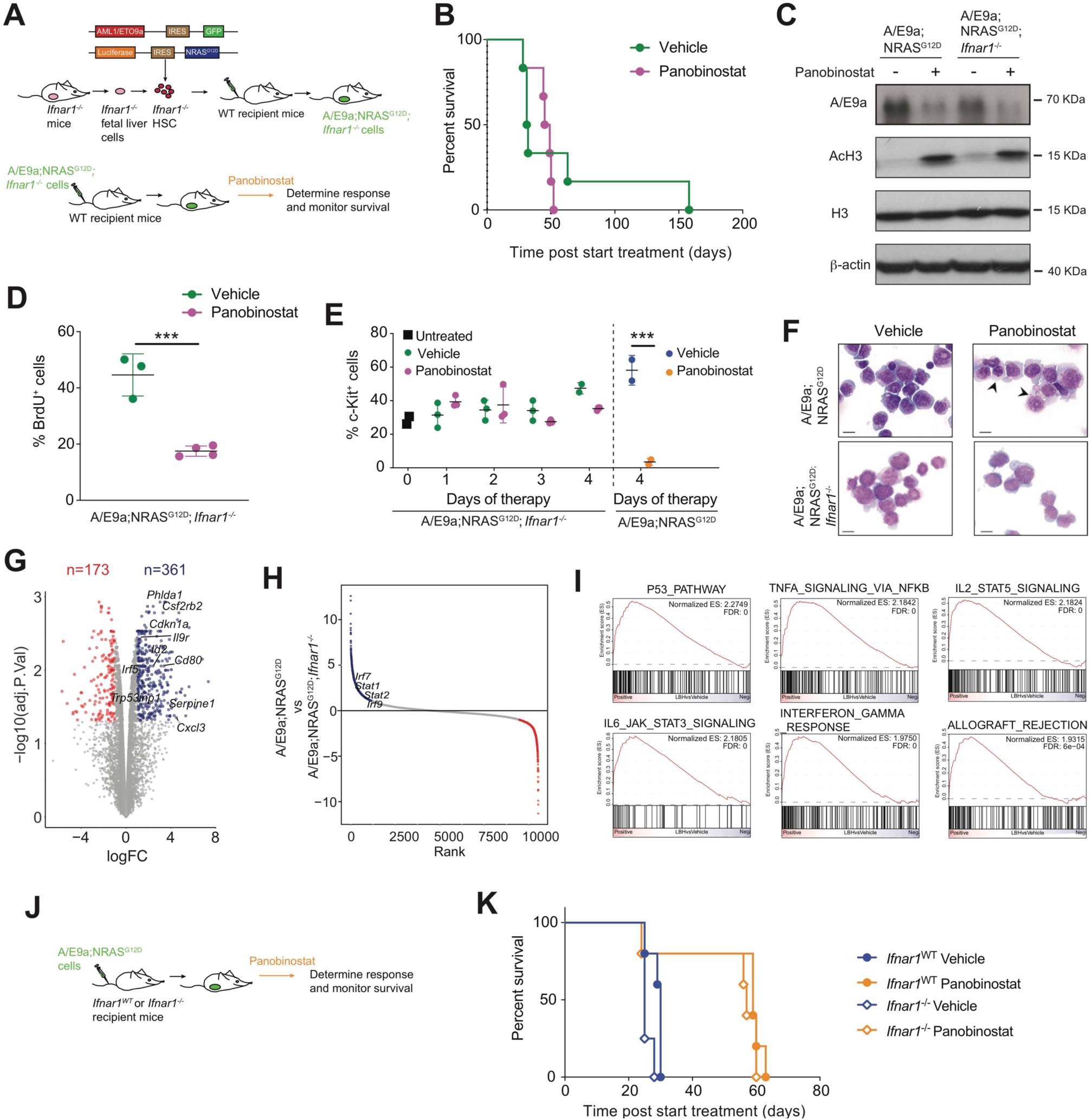
Expression of IFNAR on tumor cells is critical for the therapeutic efficacy of panobinostat. (**A**) Schematic representation of the generation of A/E9a;NRAS^G12D^;*Ifnar1*^-/-^ leukemias in wild type (WT) recipient mice. (**B**) Kaplan-Meier survival curves of mice bearing A/E9a;NRAS^G12D^;*Ifnar1*^-/-^ driven leukemias treated with either vehicle or panobinostat (n=6 mice/group). (**C**) Representative immunoblot of WT (A/E9a;NRAS^G12D^) or IFNAR-deficient (A/E9a;NRAS^G12D^;*Ifnar1*^-/-^) cells treated with either vehicle of panobinostat for 24 hours *in vitro* for the AML1-ETO9a (AE9a) fusion protein; acetylated histone 3 (AcH3); total histone 3 (H3) and β-actin as loading control (n=3). (**D**) Cell cycle analysis of A/E9a;NRAS^G12D^;*Ifnar1*^-/-^ leukemia cells within the bone marrow of mice treated with vehicle or panobinostat. The percentage of cells in S-phase (BrdU^+^ cells) was determined by flow cytometry (n=3-4; \*\*\**P*<0.001). (**E**) Flow cytometry analysis of the cell surface expression of c-Kit on A/E9a;NRAS^G12D^;*Ifnar1*^-/-^ tumor cells isolated from the bone marrow of mice treated for 4 days with vehicle or panobinostat. The differential expression of c-Kit on WT (A/E9a;NRAS^G12D^) tumor cells at day 4 is shown for comparison (n=3 mice/ timepoint; \*\*\**P*<0.001). (**F**) Representative images of May-Grünwald/Giemsa-stained WT (A/E9a;NRAS^G12D^) or IFNAR-deficient (A/E9a;NRAS^G12D^;*Ifnar1*^-/-^) tumor cells isolated from the bone marrow of mice treated with vehicle or panobinostat for 4 days. Arrows indicate condensation of nuclear chromatin and appearance of azurophilic cytoplasmic granules, indicating myeloid differentiation (600x magnification; scale bar=10μm; n=3). (**G**) Volcano plot showing DEG (logFC>1; adj.Pval<0.05) between IFNAR-deficient (A/E9a;NRAS^G12D^;*Ifnar1*^-/-^) leukemia cells isolated from panobinostat-treated and vehicle-treated leukemia-bearing mice. (**H**) Comparison of DEG between panobinostat-treated and vehicle-treated WT (A/E9a;NRAS^G12D^) and IFNAR-deficient (A/E9a;NRAS^G12D^;*Ifnar1*^-/-^) tumor cells. (**I**) GSEA of upregulated genes showed in (G). (**J**) Schematic representation of the generation of WT A/E9a;NRAS^G12D^ leukemias in *Ifnar1*^WT^ or *Ifnar1*^-/-^ recipient mice. (**K**) Kaplan-Meier survival curves of *Ifnar1*^WT^ or *Ifnar1*^-/-^ mice bearing WT A/E9a;NRAS^G12D^-driven leukemias treated with vehicle or panobinostat (n=5 mice/group).

Given that signaling through IFNAR expressed on A/E9a;NRAS^G12D^ leukemias was important for a prolonged therapeutic response to panobinostat *in vivo*, the role of type I IFN signaling in the host cells in this context was determined. Wild type C57BL/6 (*Ifnar1*^WT^) and *Ifnar1*^-/-^ mice on a C57BL/6 background (*Ifnar1*^-/-^) were transplanted with A/E9a;NRAS^G12D^ leukemias and treated with vehicle or panobinostat (**Fig. 2J**). The therapeutic effect of panobinostat against A/E9a;NRAS^G12D^ leukemias was equivalently strong in mice with (*Ifnar1*^WT^) or without (*Ifnar1*^-/-^) a host type I IFN response (**Fig. 2K** and **Suppl Fig. S2D**). These results highlight the importance of tumor-intrinsic type I IFN signaling for panobinostat-mediated differentiation and therapeutic efficacy.

### Type I IFN is produced by pDCs within the tumor microenvironment in response to panobinostat

Treatment of cancer cells with certain epigenetic drugs such as DNA methyltransferase (DNMT) inhibitors can induce a molecular and biological response that mimics anti-viral responses through reactivation of silenced endogenous retroviruses (ERVs), resulting in production of type I IFN by the tumor cells (19-21). Given that A/E9a;NRAS^G12D^ cells exposed to panobinostat displayed a type I IFN transcriptional response, we determined if IFN was being produced by the treated leukemias. No increase in IFNα or IFNβ transcription in A/E9a;NRAS^G12D^ cells was detected in response to panobinostat by genome-wide analysis using microarrays (FC>1.5, Pval<0.05; **Suppl Fig. S3A**) or gene-specific qRT-PCR (**Suppl Fig. S3B**). In the absence of evidence for production of type I IFN by the A/E9a;NRAS^G12D^ leukemias in response to panobinostat, we sought to identify the source. Dendritic cells (DCs) are capable of secreting IFNα and IFNβ in response to infection (22). These cells are phenotypically divided into pDCs (CD11c^int^Siglec-H^+^), which produce high amount of type I IFN and other cytokines in response to pathogens (23), and conventional DC (cDCs, CD11c^high^Siglec-H^-^), that activate T cells and can further be divided into cDC1 and cDC2, depending on their function and developmental origin (24).

To simultaneously examine the composition and transcriptional responses of all DC subsets, single cell CITE-seq and the related cell hashing method (25,26) was performed on DCs isolated from vehicle- or panobinostat-treated mice bearing A/E9a;NRAS^G12D^ leukemias (**Fig. 3A** and **Suppl Fig. S4A**). Concurrent experiments were performed on wild type (leukemia-free) mice to obtain baseline DC transcriptome information and content levels. Transcriptomes of single cells were generated and hashtag oligonucleotide (HTO) classification allowed the identification of the different samples (**Suppl Fig. S4B** and **Suppl Table 3**). Unsupervised clustering defined 13 distinct clusters (**Suppl Fig. S4C**), which were assigned to the different DC subpopulations previously described (24,27) by combining the transcriptional profile (**Suppl Fig. S4D-E and Suppl Table 4**) with detection of cell surface markers CD11c and Siglec-H using antibody-derived tags (ADT) (**Suppl Fig. S4F**). pDCs, cDC1s, cDC2s, CCR7^high^ DCs and monocyte-derived DC (moDCs) were identified in samples from leukemia-free and leukemia-bearing C57BL/6 mice visualized using UMAP (28) (**Fig. 3B**). Calculation of the relative proportion of each DC subset identified an expansion of the pDC compartment in leukemia-bearing mice, with 40% pDCs in vehicle-treated mice transplanted with A/E9a;NRAS^G12D^ cells compared to 12% pDCs in vehicle-treated leukemia-free mice (**Fig. 3C**). Interestingly, the DC composition in panobinostat-treated A/E9a;NRAS^G12D^ mice was restored to approximate that in vehicle-treated leukemia-free mice through the reduction on pDC numbers (**Fig. 3C**). We observed a similar trend in panobinostat-treated leukemia-free mice compared to vehicle-treated animals (**Suppl Fig. S4G**). The AUCell algorithm (29) identified that pDCs, CCR7^high^ DC and to a lesser extent cDCs, were enriched in genes from the REACTOME IFNalpha/beta signaling, which include *Ifn*α and *Ifn*β and IFN-stimulated genes (**Fig. 3D**). Treatment of mice bearing A/E9a;NRAS^G12D^ leukemias with panobinostat resulted in an increase in the type I IFN signature in pDCs and CCR7^high^ DCs populations (**Fig. 3E**). To confirm which cell type was producing type I IFN in response to panobinostat, pDCs and cDCs were isolated from vehicle- and panobinostat-treated A/E9a;NRAS^G12D^-bearing mice. Panobinostat treatment induced transcription of both *Ifn*α or *Ifn*β genes in pDCs from leukemia-bearing mice (**Fig. 3F**) In contrast, cDCs from leukemia-bearing mice did not upregulate type I IFN in response to panobinostat (**Fig. 3G**). The ability of panobinostat to induce type I IFN production from pDCs was further demonstrated using *ex vivo* pDCs from IFNβ/YFP reporter mice that had been co-treated with toll like receptor agonists poly I:C or CpG (**Suppl Fig. 5A)**. Induction of the IFNβ/YFP reporter gene was detected as early as 12 hr after incubation with panobinostat indicating that HDAC inhibition can induce *Ifn*β mRNA expression in isolated pDCs.

**FIGURE 3:**
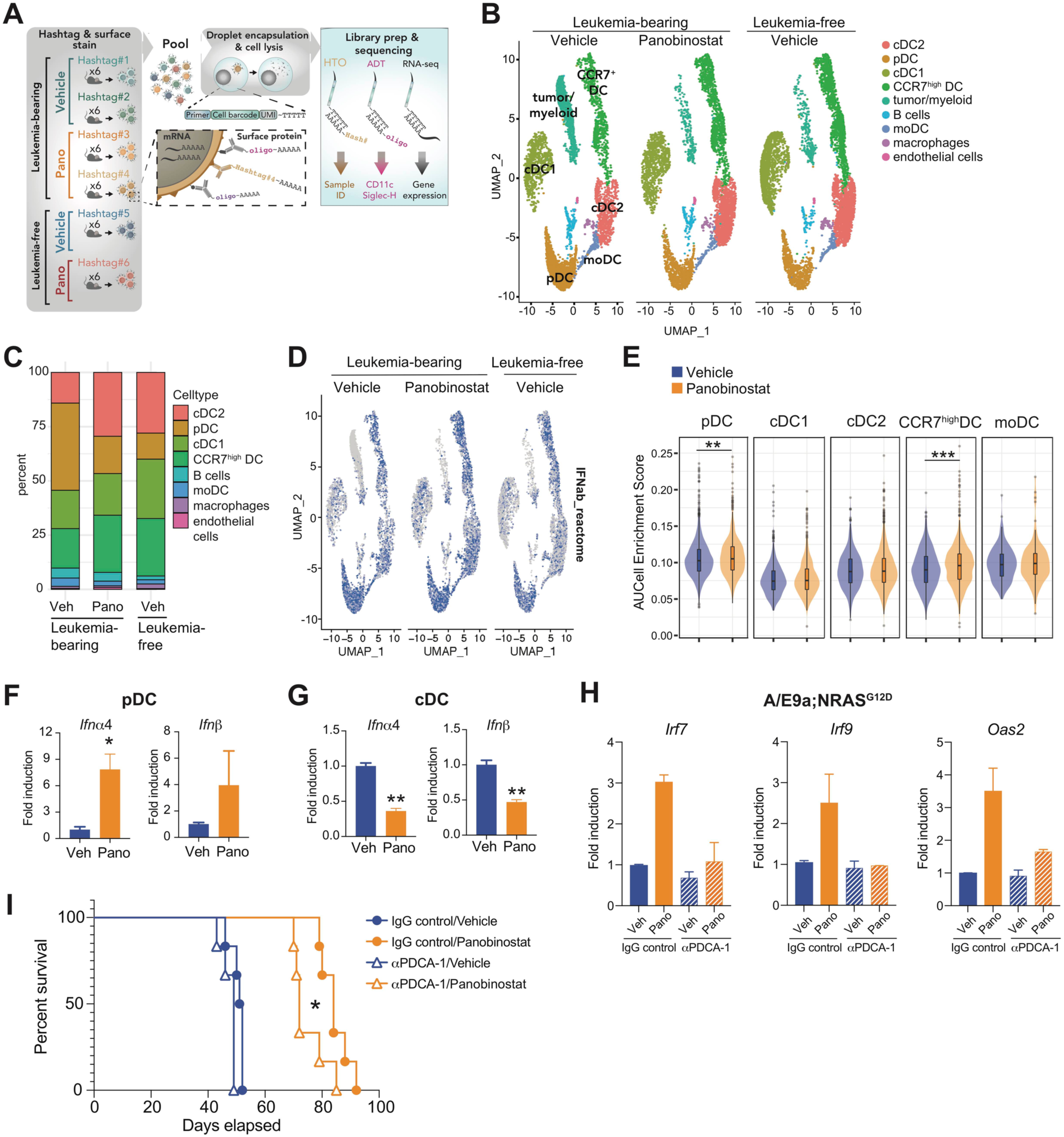
Type I IFN is produced by pDC within the tumor microenvironment in response to panobinostat. (**A**) Experimental set up for CITE-seq and Cell Hashing on DC isolated from A/E9a;NRAS^G12D^ leukemia-bearing or leukemia-free mice treated with vehicle or panobinostat for 2 days. (**B**) Annotated cell clusters based on RNA expression. (**C**) Relative frequency of individual clusters, excluding tumor/myeloid cells. (**D**) Activity of type I IFN signature on single-cell clusters. (**E**) AUCell enrichment score of type I IFN signature in DC subpopulations (\*\**P*<0.01, \*\*\**P*<0.001). (**F-G**) qRT-PCR of *Ifn*α*4* and *Ifn*β transcripts in pDC (E) and cDC (F) isolated from the spleen of mice bearing A/E9a;NRAS^G12D^-driven leukemias treated with either vehicle or panobinostat following 5 days of therapy (\**P*<0.05, \*\**P*<0.01). (**H**) qRT-PCR of *Irf7* transcript in A/E9a;NRAS^G12D^ tumor cells isolated from the spleen of mice receiving IgG control or anti-PDCA-1 and treated with either vehicle or panobinostat following 3 days of therapy (\*\**P*<0.01). (**I**) Kaplan-Meier survival curves of mice bearing A/E9a;NRAS^G12D^-driven leukemias receiving IgG control or anti-PDCA-1 and treated with either vehicle or panobinostat (n=6 mice/group; \**P*<0.05).

**FIGURE 4:**
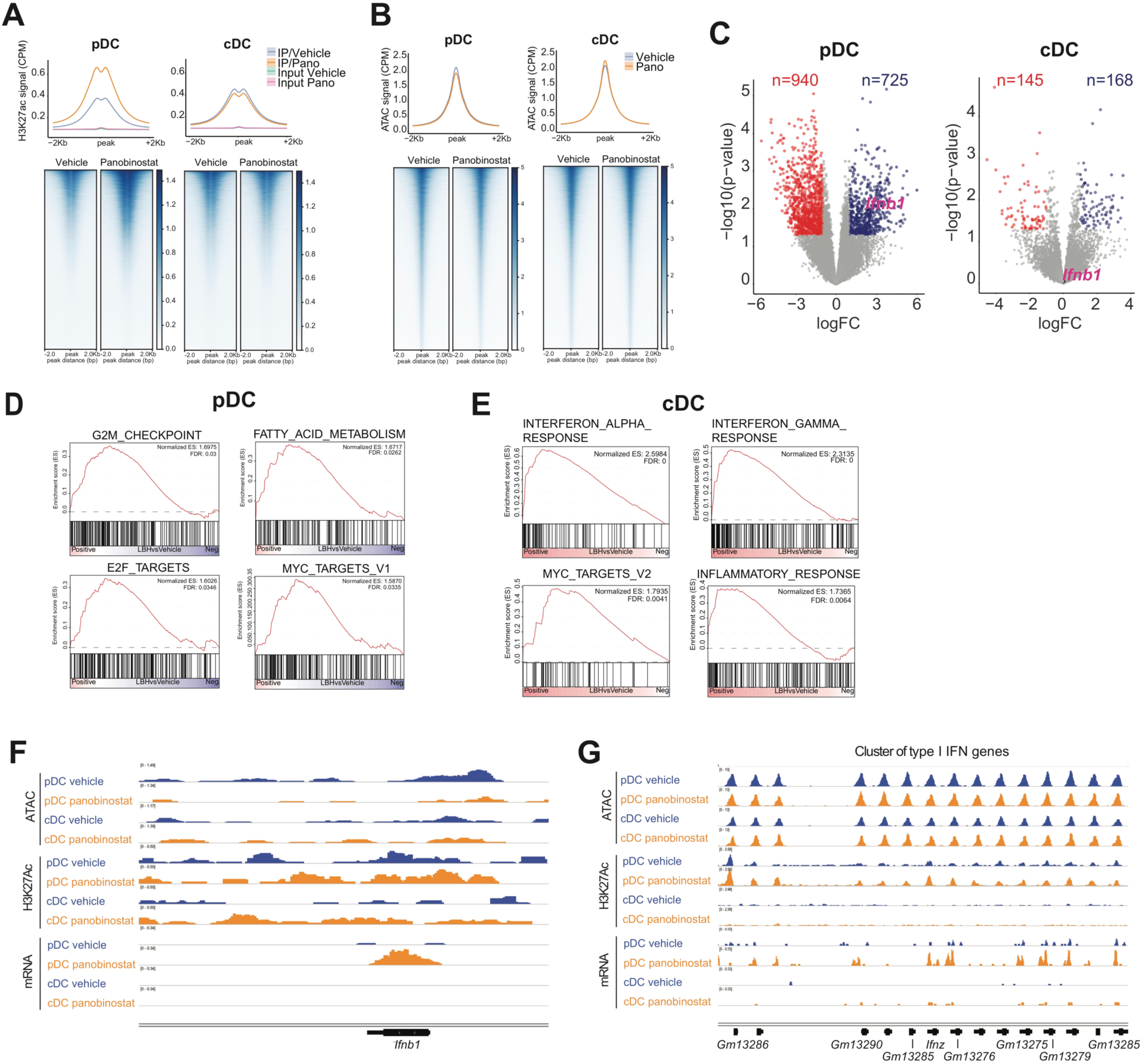
Panobinostat induces transcriptional activation of type I IFN genes in pDC through increased histone acetylation. (**A**) H3K27Ac ChIP-seq on pDC and cDC isolated from A/E9a;NRAS^G12D^ leukemia-bearing mice treated with vehicle or panobinostat for 2 days. (**B**) ATAC-seq on pDC and cDC isolated from A/E9a;NRAS^G12D^ leukemia-bearing mice treated with vehicle or panobinostat for 2 days. (**C**) Volcano plot showing DEG (logFC>1; Pval<0.05) between panobinostat-treated and vehicle-treated pDC (left) and cDC (right) isolated from A/E9a;NRAS^G12D^ leukemia-bearing mice. (**D**) GSEA of upregulated genes in pDC shown in (C). (**E**) GSEA of upregulated genes in cDC showed in (C). (**F-G**) Read-density tracks of normalized ATAC-seq, H3K27Ac ChIP-seq and RNA-seq at the *Ifnβ1* locus (F) and *Ifn*α loci (G) in pDC and cDC isolated from A/E9a;NRAS^G12D^ leukemia-bearing mice treated with vehicle or panobinostat for 2 days.

**FIGURE 5:**
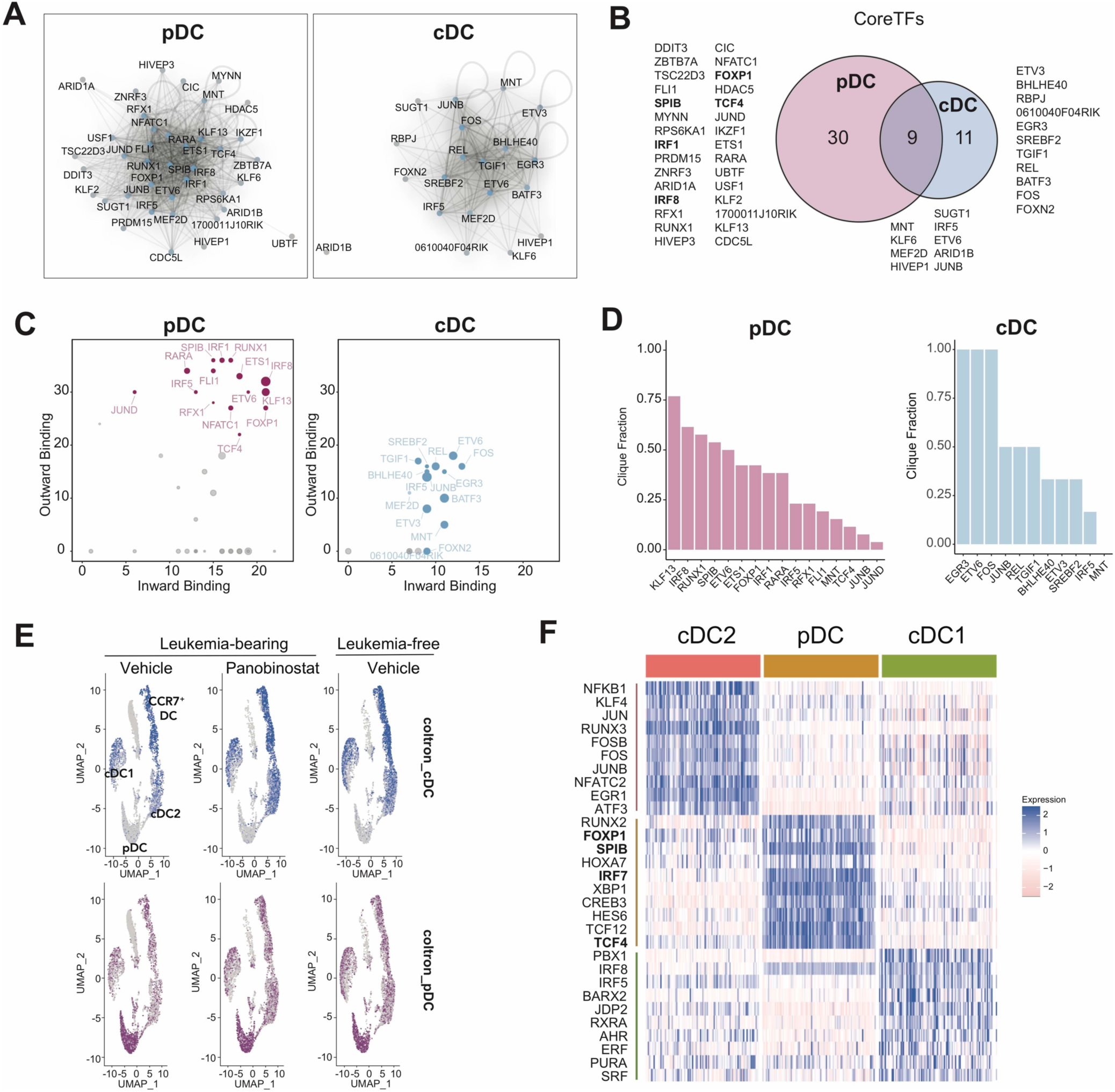
pDC depend on IRFs, the type I IFN master regulators. (**A**) CRC in pDC and cDC. (**B**) Venn diagram showing core TFs in pDC and cDC. (**C**) Inward/Outward binding plots of pDC and cDC, with highly connected core TFs highlighted. (**D**) Clique fraction for highly connected core TFs in pDC and cDC. (**E**) Activity of COLTRON-derived signatures for pDC and cDC on single-cell clusters. (**F**) Core TFs defining cell identity for pDC, cDC1 and cDC2 based on SCENIC analysis on single-cell data.

To investigate the functional role of pDC in mediating the anti-tumor response to panobinostat, an anti-PDCA1 antibody was used to deplete pDCs *in vivo* (**Suppl Fig. 5B**). Mice bearing A/E9a;NRAS^G12D^ leukemias were injected with a control antibody or pDC-depleting antibody and these mice were treated with vehicle or panobinostat for 3 days before leukemias were harvested. As shown in **figure 3H**, upregulation of type I IFN-response genes, such as *Irf7, Irf9* and *Oas2*, in A/E9a;NRAS^G12D^ cells in response to panobinostat was almost completely abrogated in the absence of pDCs. Importantly, depletion of pDCs from mice transplanted with A/E9a;NRAS^G12D^ cells resulted in a significant reduction in therapeutic efficacy mediated by panobinostat with a lower median survival of 72 days in mice receiving anti-PDCA1 compared to 84 days for IgG control mice (**Fig. 3I**). Taken together these data reveal that panobinostat induces type I IFN production in pDCs and this is important for the IFN response observed in A/E9a;NRAS^G12D^ cells as well as for the anti-leukemia effects of panobinostat.

### Panobinostat induces transcriptional activation of type I IFN genes in pDC through increased histone acetylation

The production of type I IFN in response to epigenetic drugs has been proposed to be triggered by the formation of double-stranded RNA (dsRNA) derived from ERVs, which once in the cytoplasm are sensed by pattern recognition receptors and lead to IRF7 activation through mitochondrial antiviral-signaling protein (MAVS) (21). Accordingly, the effect of panobinostat on dsRNA accumulation in pDCs was assessed. There was no detectable dsRNA in the cytoplasm of Flt3L-derived pDCs incubated with panobinostat or vehicle (**Suppl Fig. 6A**), suggesting an alternative mechanism for the induction of type I IFN in panobinostat-treated pDCs. A previous study showed that HDAC inhibition in a mouse fibroblast cell line can result in de-repression of the *Ifn*β promoter, leading to enhanced production of endogenous IFN (30). To determine if exposure of pDCs and cDCs to panobinostat resulted in increased acetylation at the regulatory regions controlling the expression of IFN genes chromatin immunoprecipation followed by sequencing (ChIP-seq) studies were performed probing for H3K27 acetylation (H3K27Ac). A genome-wide increase in H3K27Ac levels was observed in pDCs harvested from panobinostat-treated mice bearing A/E9a;NRAS^G12D^ leukemias compared to vehicle-treated mice. Interestingly, analysis of cDCs harvested from the same microenvironment showed a slight decrease in H3K27Ac following exposure to panobinostat (**Fig. 4A**). In addition, transposase-accessible chromatin by sequencing (ATAC-seq) assays (31) were performed using pDCs and cDCs harvested from transplanted mice to correlate potential panobinostat-induced changes in chromatin accessibility with transcriptional activity. Interestingly there were few significant genome-wide changes in accessible regions after panobinostat treatment in pDCs or cDCs (**Fig. 4B**).

**FIGURE 6:**
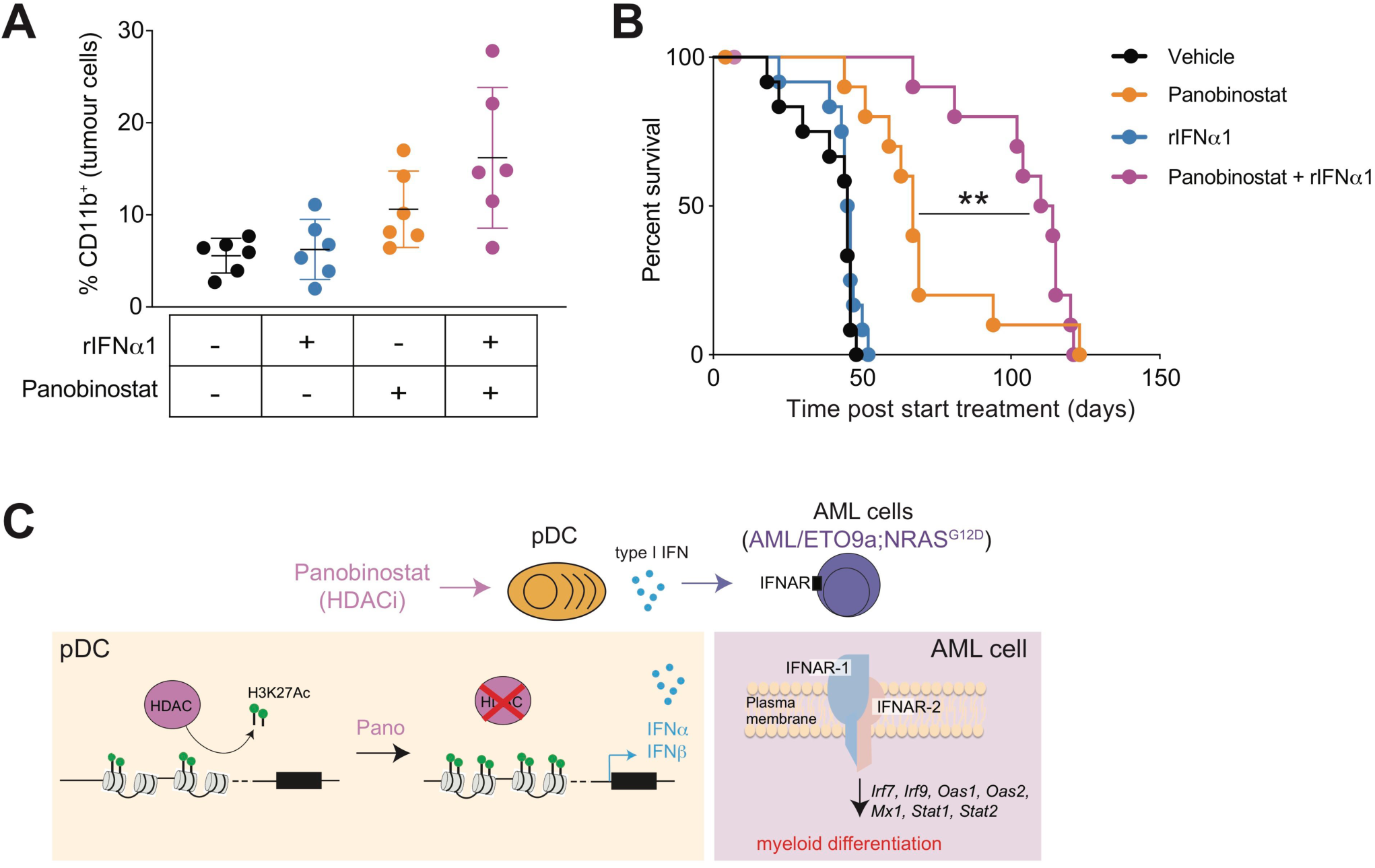
Combining panobinostat and type I IFN enhances myeloid differentiation and improves therapeutic efficacy in AML. (**A**) Flow cytometric analysis of the cell surface expression of CD11b on A/E9a;NRAS^G12D^ tumor cells isolated from the bone marrow of mice treated for 4 days with vehicle, rIFNα1, panobinostat, or a combination of rIFNα1+panobinostat (n=3). (**B**) Kaplan-Meier survival curves of mice bearing A/E9a;NRAS^G12D^-driven leukemias treated with vehicle, rIFNα1, panobinostat or the combination of rIFNα1+panobinostat (n=11-12 mice/group; \*\**P*<0.005).

Analysis of transcriptional changes mediated by panobinostat in pDCs and cDCs isolated from leukemia-bearing mice, using low input bulk RNA-seq (**Suppl Fig. 6B**) demonstrated 1665 DEG in pDCs, and only 323 DEG cDCs (**Fig. 4C** and **Suppl Tables 5 and 6**). Importantly, an increase in *Ifnβ1* transcription was detected in pDCs but not cDCs following exposure to panobinostat (**Fig. 4C**), confirming the results obtained by RT-PCR (**Fig. 3E**). GSEA and GO analysis on the 725 genes upregulated in pDCs exposed to panobinostat found enrichment in G2/M checkpoint pathway, E2F targets and metabolic pathways (**Fig. 4D** and **Suppl Fig. 6C**). Parallel analysis on the 168 genes upregulated by panobinostat in cDCs showed enrichment in pathways induced by IFNα and IFNβ, as well as IFNγ response (**Fig. 4E** and **Suppl Fig. 6D**), similar to A/E9a;NRAS^G12D^ tumor cells (**Fig. 1G**). These results suggest that pDCs in the tumor microenvironment are activated by panobinostat to produce type I IFN, which then signals through IFNAR on cDCs and A/E9a;NRAS^G12D^ leukemia cells.

To definitely determine whether panobinostat resulted in acetylation of the accessible chromatin regions of type I IFN genes in pDCs, leading to their transcriptional activation, the H3K27Ac ChIP-, ATAC- and bulk RNA-seq profiles for type I IFN genes were overlapped in pDCs, and cDCs as comparison. A clear increase in H3K27Ac at the *Ifnβ1* gene (**Fig. 4F**) and more generally in the cluster of type I IFN genes (**Fig. 4G**) was detectable in pDCs harvested from panobinostat-treated leukemia-bearing mice (orange profiles), compared to pDCs from vehicle-treated mice (blue profiles). Enhanced H3K27ac at type I IFN gene loci observed in pDC exposed to panobinostat correlated with increased transcription of the corresponding genes based on mRNA reads, and overlapped with open chromatin regions (**Fig. 4F and 4G**). In contrast, analysis of cDC showed that panobinostat treatment barely affected H3K27Ac levels at type I IFN genes and little if any detectable increase in transcription of those genes was observed (**Fig. 4F and 4G**). Interestingly, low level H3K27Ac marks and mRNA reads were observed in pDCs, but not cDCs, from vehicle-treated mice (**Fig. 4F and 4G**), suggesting that pDCs are transcriptionally primed to produce type I IFN.

### pDC identity is linked to IRFs, the master regulators of type I IFN

The data detailed above indicated that despite both pDCs and cDCs demonstrating open chromatin regions around type I IFN genes, only pDC were putatively primed to produce type I IFN following exposure to panobinostat (**Fig. 4F and 4G**). It is possible that that these differences were due to the expression and/or activity of core transcription factors (TFs) distinct for each cell type. To test this hypothesis, the information derived from H3K27Ac ChIP-seq studies on vehicle-treated pDCs and cDCs were used to define the enhancer landscape of each cell type. This allows the identification of super enhancers (SE), broad open regions of chromatin with high levels of H3K27Ac that facilitate binding of the core TFs that control the transcriptional program of a given cell (32,33). SE for pDCs and cDCs were defined by stitching H3K27ac peaks within 12.5Kb in each cell type and ranking them by H3K27Ac signal (**Suppl Fig. 7A**). Subsequent identification of SE target genes revealed 584 in pDCs and 299 in cDCs, with only 16% shared between both populations (**Suppl Fig. 7B and Suppl Table 7**). TFs associated with SEs were identified, and open regions within such SEs were used to identify TF motifs and create TF-binding networks (i.e. core regulatory circuitries, CRC) for pDCs and cDCsusingCOLTRON (https://pypi.python.org/pypi/coltron) (**Fig. 5A**). Importantly, when the CRC of pDCs and cDCs were overlapped only 9 shared core TFs were identified(**Fig.5B**).Overall TF connectivity in the network was visualized by calculating the number of TF motifs within the SE of a node TF (inward binding) and the number of TF associated SEs bound by a node TF (outward binding) (32-34) (**Fig. 5C**). Among the most connected core TFs in pDCs were TCF-4, IRF1 and IRF8, previously described as important for pDC development and function(35,36),andknownfactors associated with B-cell lineage, including SPIB and FOXP1 (**Fig. 5C**). This was further confirmed when the clique fraction of the core TFs (see Methods) were calculated, and SPIB, FOXP1, IRF8 and IRF1 were TFs identified to dominate clique membership in pDCs (**Fig. 5D**). Importantly, both IRF1 and IRF8 have been shown to be involved in type I IFN-mediated responses (36,37). As an orthogonal approach, single cell RNA-seq data from figure 3 was utilized. AUCell analysis confirmed that the pDCs and cDCs signatures defined by COLTRON were active in pDCs and cDCs subsets respectively, defined at the single-cell level (**Fig. 5E**), thus validating the regulatory networks established using COLTRON.

Application of the SCENIC algorithm, which infers regulatory networks based on predicted TF activity (29) was used to explore in more detail the CRC of pDCs, cDC1s and cDC2s. Clustering using SCENIC (**Suppl Fig. 7C**) identified the same cell types as clustering based only in the transcriptional profile (**Fig. 3B**). The TFs driving the identity of each cell type were then identified (**Fig. 5F** and **Suppl Table 8**) revealing TCF4, FOXP1, SPIB; and also IRF7, the master regulator of type I IFN-dependent response (38), in pDCs which were not present in cDC1 or cDC2 clusters (**Fig. 5F**). This analysis also showed IRF8 as a SCENIC marker for pDC and cDC1 (**Fig. 5F**). These results indicate that the unique capacity of pDC to produce IFN in response to panobinostat likely relies on their dependency for cell identity on IRFs, which are known to be essential for the transcriptional activation of type I IFN genes. In addition, these data support a different origin for pDC and cDC, with pDC closer to the B-cell lineage, as has been recently proposed (39).

### Combining panobinostat and recombinant type I IFN enhances myeloid differentiation and improves therapeutic efficacy in AML

Given the results linking activation of type I IFN signaling to panobinostat mediated differentiation of A/E9a;NRAS^G12D^ cells and the therapeutic efficacy of panobinostat in this setting, the combination of panobinostat and recombinant IFNα1 (rIFNα1) was tested in leukemia-bearing mice. To determine the effect of this combination on myeloid differentiation, CD11b expression on the tumor cells following 4 days of therapy was assessed. In contrast to single agent panobinostat treatment, rIFNα1 alone had no effect on the myeloid differentiation status of these cells. The combination of rIFNα1 with panobinostat resulted in an increase CD11b expression that did not reach statistical significance (**Fig. 6A**). More importantly, while rIFNα1 had no therapeutic benefit in mice bearing A/E9a;NRAS^G12D^ leukemias, the combination of rIFNα1 and panobinostat resulted in a significant increase in the survival of tumor-bearing compared to those mice treated with panobinostat alone. The median survival benefit of panobinostat+rIFNα1 over vehicle was 79.5 days, compared to 32.5 days for single agent panobinostat treatment over vehicle (**Fig. 6B**). These results highlight the therapeutic potential of combining HDACi and type I IFN as a new approach to treat t(8;21) leukemia.

## DISCUSSION

Since AML is characterised by deregulation of epigenetic modifications, which contribute to the pathogenesis of the disease, targeting the epigenome through the pharmacological inhibition of epigenetic modifiers has been the main focus of many preclinical and clinical studies in leukemia during the last two decades (40). This previous work has clearly demonstrated that epigenetic drugs, such as HDACi and DNMT inhibitors (DNMTi), have multiple mechanisms of action that result in loss of leukemia cell proliferation and survival, including through the activation of the immune system (41). DNMTi have been shown to induce a type I IFN response, activating the IFNAR-mediated pathway, which can contribute to the anti-tumor effects of these anti-cancer agents (19,20). However, these previous studies with epigenetic therapies focused on the pathways leading to type I IFN production by cancer cells, and the direct impact of these compounds within the immune compartment was not characterized. In this regard, recent evidence strongly indicates that epigenetic agents can also regulate the function of immune cells in the tumor microenvironment and, therefore, they have the capacity to modulate the anti-tumor immune response (40,41). Here we demonstrate that the panobinostat-mediated induction of type I IFN by host pDCs is required for the full therapeutic benefit of HDACi in pre-clinical models of t(8;21) AML (**Fig. 6C**). This novel immunomodulatory mechanism initiated by the effect of an epigenetic drug on immune cells, rather than the tumor, highlights the importance of understanding the global effects triggered by epigenetic therapies to maximize their therapeutic potential in the clinical setting.

Mechanistically, the IFN-inducing activity of panobinostat was specifically confined to pDCs and the induction of type I IFN genes in these cells was concomitant with panobinostat-mediated hyperacetylation within the *Ifn* loci. The activation of type I IFN production occurred at the transcriptional level through increased H3K27Ac and, therefore, is different from the viral mimicry mechanism proposed for other epigenetic inhibitors, which induce the expression of ERV-derived dsRNA that is detected by intracellular sensors, leading to type I IFN transcription (21). Interestingly, only pDCs but not cDCs responded to HDAC inhibition by upregulating type I IFN transcription. We found that pDCs were in a pre-activated state before treatment, with relatively high basal histone acetylation levels in type I IFN genes. This basal H3K27Ac level is probably influenced by the transcriptional dependency on IRFs that we have shown for pDCs and drives cell identity of this population. To our knowledge, the present study is the first to describe the CRC for pDC, cDC1 and cDC2, identifying the TF interactions that regulate the transcriptional program in each cell type.

We and others previously demonstrated that the use of HDACi in mouse models of t(8;21) AML resulted in leukemia cell differentiation, cell cycle arrest and apoptosis (7,42-45). The results provided herein unveil that panobinostat also triggers a very strong activation of the type I interferon receptor (IFNAR) signaling pathway in leukemia cells, which is essential for myeloid differentiation of the tumor cells and the resultant therapeutic outcome (**Fig. 6C**). These results are in line with previous work demonstrating that activation of the type I IFN pathway in leukemia cells has anti-proliferative and pro-apoptotic effects, and promotes differentiation, resulting in abrogation of AML development (16,46). Type I IFN therapy has previously been tested as a treatment option in AML, although these clinical trials have shown only limited success (47). One plausible explanation is the high toxicity and low stability of the IFN preparations used in those studies. To overcome this issue, more recent pegylated versions of recombinant type I IFN have been formulated, improving their tolerability and durability (48). Yet, it is also possible that IFN has limited anti-leukemic effect as a single agent and its clinical applicability will be improved in combination with other therapies, as our results suggest.

The use of HDACi as a single agent has resulted in moderate therapeutic efficacy in early phase clinical trials of AML (10,49). Recent studies combining HDACi with other anti-cancer agents have shown improved clinical outcomes (10,47,49). Specifically, combinations of HDACi with other anti-cancer drugs such as hypomethylating agents, all-trans retinoic acid (ATRA), chemotherapy or BET inhibitors are currently being tested in the clinic for AML patients (47,50), with panobinostat in combination with chemotherapeutic agents showing very promising results (51-53).

Another option is to combine epigenetic drugs with immunotherapy, since mounting evidence shows that modulation of the immune system through epigenetic therapies can be leveraged to design better treatments for hematological malignancies and many other types of cancer (21). In line with this, our results indicate that a regimen combining HDAC inhibition with type I IFN could significantly improve the efficacy of HDACi as anti-cancer treatments.

In summary, detailed single cell molecular analysis using integrated genomics was used to discover a novel mechanism of epigenetic regulation of immune cells, specifically pDCs, that can be harnessed to improve the therapeutic effects of HDACi in t(8;21) AML. The results of our study highlight the importance of studying the effects of systemic therapies on both the host immune cells and tumor cells and demonstrates the complex molecular interplay between cells and signaling molecules within the tumor microenvironment. We harnessed this information to derive a new therapeutic regimen that was vastly superior to single agent therapy. These results strongly suggest that combining HDAC inhibition with type I IFN would benefit leukemia patients with t(8;21) translocations.

## METHODS

### Experimental mice and materials

Wild-type C57BL/6 mice at 6-8 weeks of age were purchased from The Walter and Eliza Hall Institute of Medical Research. *Ifnar1*^-/-^ null mice (18) were a generous gift from Paul Hertzog (Monash University). Animals were maintained under specific pathogen-free conditions and used in accordance with the institutional guidelines approved by the Peter MacCallum Cancer Centre Animal Ethics Committee (Approval numbers E472, E555, E627).

Panobinostat (LBH589) was kindly provided by Novartis and prepared as a 2mg/ml solution in 5% dextrose/dH_2_O (D5W). Recombinant mouse interferon-alpha (rIFNα1), was kindly provided by Prof. Paul Hertzog (Monash University, Australia) and diluted in phosphate-buffered saline (PBS). Anti-PDCA-1 (*InVivo*MAb anti-mouse CD317 Cat# BE0311) and IgG control (*InVivo*MAb anti-keyhole limpet hemocyanin Cat# BE0090) were purchased from BioXCell. Poly(I:C) (HMW)/Lyovec was purchased from InvivoGen (tlrl-piclv). CpG 2216 (Cat# 930507) was obtained from Tib Molbiol and complexed to DOTAP liposomal transfection reagent (Roche, Cat# 11202375001) for increased type I IFN induction. Poly I:C (Cat# 27-4732-01) was purchased from GE Healthcare).

### Generation of A/E9a;NRAS^G12D^ driven leukemias

Retroviral transduction of mouse fetal liver cells (wild-type or *Ifnar1*^-/-^) was performed as previously described (7). Briefly, viral producer cells (Phoenix) were transfected with MSCV-AML1/ETO9a-IRES-GFP or MSCV-luciferase-IRES-NRAS^G12D^ vectors. Viral supernatants were mixed at a 1:1 ratio and used to transduce mouse fetal liver cells. To generate transplantable AML, 1×10^6^ total cells per recipient were injected into lethally irradiated C57BL/6 mice (2x 5.5Gy; 4h apart) via the tail vein. To prevent infections, transplanted mice were provided with water containing neomycin and polymyxin B (both from Sigma).

### Monitoring leukemia

Tumor burden in transplanted mice was assessed by whole-body bioluminescent imaging (BLI) using an IVIS100 imaging system (Perkin Elmer) and blood sampling. For BLI, mice were injected intraperitoneally with 50mg/kg D-Luciferin (Perkin Elmer), anesthetized with isoflurane, and imaged for 2min. Peripheral blood white cell counts (WCC) were measured using an Advia 120 automated hematology analyzer (Bayer Diagnostics) and percentage of GFP-positive cells was analyzed by flow cytometry. At terminal disease stage, mice were euthanized and leukemia cells were isolated from bone marrow (femur) and spleen. Single cell suspensions were prepared and cells were cryopreserved in FCS/10% DMSO. To analyze differentiation of leukemia cells, May-Grünwald Giemsa (MGG) stain was performed on cytospins of FACS sorted (GFP positive) bone marrow cells.

### *In vivo* therapy

Cryopreserved leukemia cells were thawed, washed and resuspended in PBS. Approximately 1×10^6^ cells per recipient were transplanted into sub-lethally irradiated mice (2x 3Gy; 4h apart) by tail vein injection. Mice were monitored weekly and treatment was initiated once leukemia was clearly established (corresponding to 5-20% GFP-positive cells in peripheral blood). For therapy studies, mice were treated daily with 25mg/kg panobinostat, five consecutive days per week by intraperitoneal injection for one week followed by 15mg/kg panobinostat for three weeks. Control mice received the equivalent volume of vehicle. Mortality events from advancing leukemia were recorded for the analysis of therapeutic efficacy. For short-term drug response studies, mice received 25mg/kg panobinostat, or equivalent volume of vehicle by intraperitoneal injection once daily for indicated period prior to harvesting of spleen and bone marrow. Cell suspensions were prepared and used for further analyses.

### Flow cytometry

Cell suspensions were lysed in red cell lysis buffer (150mM NH_4_Cl, 10mM KHCO_3_, 0.1mM EDTA) and washed twice in FACS staining buffer (PBS supplemented with 2% FCS and 0.02% NaN_3_). Aliquots of cells were pre-incubated with anti-CD16/CD32 (2.4G2) and stained on ice with anti-CD117 (c-Kit), anti-CD11b (Mac-1), anti-Ly6A/E (Sca1), anti-IFNAR1, anti-B220 (all from BD Biosciences), anti-CD11c, anti-SiglecH (both from Biolegend) in FACS staining buffer for at least 30min. Tumor cells were identified using GFP reporter expression. For cell proliferation analysis, BrdU (100μl of a 10mg/mL stock; Sigma) was injected intraperitoneally in to mice 16 hours prior to euthanasia. Bone marrow cells were fixed, permeabilized and stained using BrdU Flow Kit (BD Pharmingen), following the manufacturer’s instructions. For dsRNA staining, cells were fixed and permeabilized (BD Pharmingen) and stained with K1-IgG2a monoclonal antibody (SCICONS, Cat# 10020200) that recognizes dsRNA. To be detected by flow cytometry, K1-IgG2a was previously conjugated with APC using the APC conjugation kit from Innova Biosciences (Cat# 705-0030). Data was collected on a FACSCanto II flow cytometer (BD Biosciences) and analyzed using FlowJo software (Tree Star).

### Splenic dendritic cell isolation

Spleens were collected into RPMI 1640 containing 2% fetal calf serum (FCS) and dissociated with a scalpel and type 3 collagenase (Worthington) and DNAse I, grade 2 (Roche), homogenizing for 20min with a plastic Pasteur pipette. Then 0.1M EDTA was added and the sample was homogenized for another 5min. Single cell suspensions were filtered through 40μM cell strainers. Samples were enriched in dendritic cells using OptiPrep (Axis-Shield Cat# 1114542) gradient, following the manufacturer’s protocol for dendritic cell isolation from tissues. pDC and cDC populations were sorted in a BD FACSAria Fusion Cell Sorter (BD Bioscience) based on CD11c and SiglecH expression. For single cell RNA-seq, spleens were enriched in DC using OptiPrep and leukemia cells were removed through sorting, collecting GFP^-^ cells. Then DC were negatively selected using the EasySep Mouse Pan-DC Enrichment Kit (Stem Cell Technologies, Cat# 19763).

### *Ex vivo* stimulation of bone marrow cells

Femur and tibia bones from IFNβ/YFP reporter mice (54) were isolated and kept in PBS. Bone marrow was flushed with RPMI 1640 medium without FCS using a 26G needle. Single cell suspensions were lysed using Erylysis-buffer (Morphisto) and filtered through 70μM cell strainers. Cells were cultured in RPMI 1640 medium containing 1.6 nM panobinostat or vehicle control (DMSO) for 24h before 1µM CpG 2216 or 0.1 µM poly I:C was added to the culture for additional 12h or 24h. Percentage of IFNβ/YFP^+^ cells of pDCs (gated as CD11c^int^ B220^+^ CD11b^-^) was determined by flow cytometry.

### Flt3L-derived dendritic cell cultures

Bone marrow-derived Flt3L-cultured pDCs were generated as previously described (54). In short, single cell suspensions from mouse bone marrow were generated as described above. Cells were seeded at 20×10^6^/10cm petri dish (cell culture non-treated plates) in RPMI 1640 medium containing 100ng/ml recombinant Flt3L (BioXCell). At day 5, cells were collected and fresh media added. pDC were sorted at day 7 (gated as CD11c^+^Siglec-H^+^) and incubated with vehicle, panobinostat or poly I:C for 6h.

### Immunoblot

Protein lysates were prepared using cell lysis buffer (150mM NaCl, 10mM Tris-HCl [pH 7.4], 5mM EDTA, 1% Triton X-100) supplemented with PMSF and the cOmplete(tm) protease inhibitor cocktail (EDTA-free; Roche). Lysates were subjected to SDS-PAGE and transferred to PVDF membrane (Millipore). Membranes were blocked in skim milk or BSA and probed overnight with the following antibodies: Anti-AML1 (Cell Signaling Technology, Cat# 4336); anti-acetyl-histone H3 (Millipore, Cat# 06-599); anti-histone H3 (Abcam, Cat# 1791) and anti-β-actin (Sigma, Cat# AC-74). Filters were then washed, probed with either anti-mouse HRP, anti-rabbit HRP (both DakoCytomation) secondary antibodies. Signals were visualized using enhanced chemiluminescent detection reagent (Amersham).

### Quantitative RT-PCR

Total RNA from FACS sorted tumor cells (GFP^+^) was isolated using TriZol (Invitrogen) following manufacturer’s protocol. Synthesis of cDNA was performed following standard protocols using MMLV Reverse Transcriptase (Promega). Quantitative RT-PCR was performed using SYBR Green (Applied Biosystems) method in a 384-well format using the ABI Prism 7900HT (Applied Biosystems) or a 96-well format StepOne Plus (Applied Biosystems). For quantification, the C_T_ values were obtained and normalized to the C_T_ values of *Hprt* or *Actin* gene. Fold changes in expression were calculated by the 2^-ΔCT^ method. Primer sets used for gene expression analysis can be found in Supplementary Table 9.

### Microarray, RNA-sequencing and analysis

Following three days of treatment with panobinostat or vehicle, tumor cells (GFP^+^) were sorted on a BD FACSAriaII cell sorter (BD Biosciences), propidium iodide (PI) was used as a viability marker. Total RNA was prepared from three independent replicates using Trizol (Invitrogen) following manufacturer’s protocol. Samples were further purified using the RNeasy MinElute kit (Qiagen) or Direct-zol RNA Miniprep kit (Zymo Research). mRNA libraries for A/E9a;NRAS^G12D^ tumor cells were generated using TruSeq RNA Library Prep kit (Illumina) and sequenced in a HiSeq2000 sequencer SE50bp (Illumina). Microarray for A/E9a;NRAS^G12D^ tumor cells was performed using Affymetrix MoGene-1_0-st-v1. Data was analysed using Limma in R. mRNA libraries for A/E9a;NRAS^G12D^;*Ifnar1*^-/-^ tumor cells were generated using TruSeq RNA Library Prep kit (Illumina) and sequenced in a HiSeq2500 sequencer PE50bp (Illumina). mRNA libraries for DC, sorted from spleens of A/E9a;NRAS^G12D^-leukemia bearing mice after two days of treatment with panobinostat or vehicle, were generated using NEBNext Single Cell/Low Input RNA Library Preparation kit (New England Biolabs) and sequenced in a NextSeq500 sequencer SE75bp (Illumina). Sequencing files were demultiplexed (Bcl2fastq, v2.17.1.14) and QC was performed on FASTQ files using FASTQC (v0.11.6). Sequencing reads were trimmed (cutadapt v2.1) and aligned to the mm10 mouse reference genome using HISAT2 (v2.1.0). Read counting across genomic features was performed using FeatureCounts (Subread, v2.0.0) and differential gene expression analysis was performed using Voom-Limma in R (v3.42.2). Gene set enrichment analyses were performed using the Broad Institute GSEA software (55).

### ChIP-sequencing

20,000 DC, sorted from spleens of A/E9a;NRAS^G12D^-leukemia bearing mice after two days of treatment with panobinostat or vehicle, were fixed with 1% formaldehyde for 6min and sonicated using the S220 Focused-ultrasonicator (Covaris) for 8min. Samples were incubated with an antibody to H3K27Ac (Abcam Cat# ab4729) and chromatin was isolated with the True Microchip Kit (Diagenode Cat# C01010130) according to the manufacturer’s instructions. Libraries were prepared using the DNA Library Prep NEB (New England Biolabs) and sequenced in a NextSeq500 sequencer SE75bp (Illumina).

Bcl2fastq (v2.17.1.14) was used to demultiplex sequencing files and resulting Fastq output was quality checked using FASTQC (v0.11.6). Fastq read alignment to the mouse genome (mm10/GRCm38) was performed using Botwtie2 (v2.3.4.1). SAM files generated were converted to BAM file format and subsequently sorted, indexed and potential PCR duplicates were removed using Samtools (v1.9) with view, sort, index and rmdup functions, respectively. Deeptools (v3.0.0) bamCoverage function was used for BAM to BigWig file conversion using the following settings (--normalizeUsing CPM --smooth length 150 -bs 50 -e 225).

BigWig file average read density across defined genomic intervals was performed using Deeptools (v3.0.0) computeMatrix function with the following settings (reference-point --referencePoint center -- upstream 2000 --downstream 2000 -bs 50 --missingDataAsZero), where average read density was calculated +/- 2Kb of ATAC peak summits and genomic regions spanning mm10 blacklist intervals (ENCODE, accession ENCFF547MET) were removed. Resulting matrices were subsequently used to generate heatmap and average profile plots with Deeptools (v3.0.0) plotHeatmap and Rstudio (v3.6.1), respectively.

BAM file peak calling was performed MACS (v2.1.1) with the following settings (-f BAM -m 10 30 --broad --cutoff-analysis -g mm -c INPUT) and resulting BED files were filtered for regions spanning mm10 blacklist intervals (ENCODE, accession ENCFF547MET) and modified into GFF file format using Rstudio (v3.6.1) to meet input criteria for ROSE2 (v1.0.5).

### ATAC-sequencing

15,000 DC, sorted from spleens of A/E9a;NRAS^G12D^-leukemia bearing mice after two days of treatment with panobinostat or vehicle, were lysed in cold ATAC-seq cell lysis buffer and exposed to Tn5 transposase (Nextera) for 30min at 37°C. Transposed chromatin was amplified using customized Nextera PCR Primer 1 and Primer 2 (barcode) (31) and HotStart KAPA ReadyMix (Roche). Libraries were quantified by Qubit using the DS DNA HS kit (Thermo Fisher Scientific), QC validated using the Tape Station DNA High Sensitivity D1K (Agilent), size-selected using Pippen Prep 1.5 % Agarose Gel cassettes (Sage Science) and sequenced in a NextSeq500 sequencer SE75bp (Illumina).

Similar to ChIP-seq analysis, Bcl2fastq (v2.17.1.14) was used to demultiplex sequencing files and resulting Fastq output was quality checked using FASTQC (v0.11.6). Fastq read alignment to the mouse genome (mm10/GRCm38) was performed using Botwtie2 (v2.3.4.1). SAM files generated were converted to BAM file format and subsequently sorted, indexed and potential PCR duplicates were removed using Samtools (v1.9) with view, sort, index and rmdup functions, respectively. Deeptools (v3.0.0) bamCoverage function was used for BAM to BigWig file conversion using the following settings (--normalizeUsing CPM --smooth length 150 -bs 50 -e 225).

BAM file peak calling was performed MACS (v2.1.1) with the following settings (-g mm -f BAM --call-summits --nomodel --extsize 100). BigWig file average read density across defined genomic intervals was performed using Deeptools (v3.0.0) computeMatrix function with the following settings (reference-point --referencePoint center --upstream 2000 --downstream 2000 -bs 50) where average read density was calculated +/- 2Kb of ATAC peak summits.

### Single cell sequencing methods: Ab-staining, capture of cells in GEMs, barcoding of transcriptomes and library preparation

DCs -isolated from spleens through Optiprep gradient, followed by sorting of GFP^-^ cells and magnetic negative selection-were incubated with Cell Hashing HTO-conjugated antibodies (Biolegend, TotalSeq(tm) anti-mouse Hashtag reagents Cat# A0301-A0306) and CITE-seq antibodies (Biolegend, TotalSeq(tm)-A0106 anti-mouse CD11c and A0119 anti-mouse Siglec H), as described previously (25,26). Cells were washed 3 times with FACS buffer and samples pooled at an equal concentration of 1500 cells/μl. Pooled DC samples were loaded onto the 10x Chromium instrument (10x Genomics, Pleasanton, CA, USA) to generate single-cell Gel Beads-in-Emulsion (GEMs) and capture/barcode cells. All samples followed the 10x Genomics Single Cell 3′ v3 according to the manufacturer’s instructions up until the cDNA amplification step (10x Genomics, USA). Two picomoles of HTO and ADT additive oligonucleotides were spiked into the cDNA amplification PCR, and cDNA was amplified according to the 10x Single Cell 3′ v3 protocol (10x Genomics, USA). Following cDNA amplification, 0.6X SPRI was used to separate the large cDNA fraction derived from cellular mRNAs (retained on beads) from the ADT- and Cell Hashtag (HTO)-containing fraction (in supernatant). The cDNA fraction was processed according to the 10x Genomics Single Cell 3′ v3 protocol to generate the transcriptome library; indexing was done using Chromium i7 Multiplex Kit. An additional 1.4X reaction volume of SPRI beads was added to the ADT/HTO fraction to bring the ratio up to 2.0X. The beads were washed with 80% ethanol, eluted in water, and an additional round of 2.0X SPRI performed to remove excess single-stranded oligonucleotides from cDNA amplification. After final elution, separate PCRs were set up to generate the CITE-seq ADT library (SI-PCR and RPI-x primers) and the HTO library (SI-PCR and D7xx_s).

### Single cell RNA-sequencing analysis

10X Genomics’ Cell Ranger Software (v3.1.0) was used to align reads to the mm10/GRCm38 reference genome, demultiplex cellular barcodes and quantify unique molecular identifiers (UMIs) and antibody capture (Hashtag oligos (HTOs) and Antibody Derived Tags (ADTs)). Inter-sample doublets and intra-sample doublets were removed using Seurat’s HTODemux function and the Scrublet python package (v0.2.1) (56), respectively. In more detail, counts with more than one median absolute derivation (MAD) above the median Scrublet score were retained.

Single-Cell Regulatory Network Inference and Clustering (SCENIC) method (29) was used to estimate relative transcription factor activity in each cell. Calculation of Area-under-curve (AUC) scores for transcription factors was performed using inferred using gene regulatory networks with the pyscenic python package (v3.6.1).

The Seurat R package (v3.1.0) (57) was used to process gene expression and antibody count matrices within R studio (v3.6.1). Seurat’s SCTransform function was subsequently used to sctransform normalise RNA transcript counts for barcodes identified as cells by Cell Ranger. Transformation of counts with no covariates in the sctranform model was first performed, then Seurat’s CellCycleScoring function with cell cycle gene sets of mouse homologues was used to calculate cell cycle phase scores. Cell cycle phase scores and the percent of raw RNA counts belonging to mitochondrial genes for each cell were used to rerun sctransform normalisation and regress out of the model.

Centred log ratio (CLR) transformation was performed to normalise ADT counts.

Sctransform scaled RNA expression values for genes with residual variance greater than 1.3 in the sctransform model were used for Principle component analysis (PCA). The top 10 principle components were used to calculate a shared-nearest-neighbours (SNN) network with the FingNeighbors function with a cosine distance metric and 50 k-nearest neighbors. Cell populations were subsequently identified using the SNN network and the FindClusters function with the Louvain algorithm and a resolution parameter of 0.6. The top 10 principle components were also used to calculate Uniform Manifold Approximation and Projection (UMAP) values using the RunUMAP function with a cosine distance metric and 50 nearest neighbors as parameters.

### Analysis of core regulatory circuitries

Putative super-enhancer regions from H3K27ac ChIP-seq peaks were identified using the Ranking Ordering of Super-Enhancer (ROSE2; v1.0.5) algorithm (58,59) with the following settings (-g MM10 -c INPUT -s 12500 -t 2000), where enhancer peaks within 12.5Kb were stitched and peaks within 2Kb of a TSS were excluded to remove promoter bias.

The Coltron (v1.0.2) algorithm was used to construct transcription factor regulatory networks using H3K27ac ChIP-seq signal (BAM file) and ROSE2 generated enhancer table to identify super-enhancer associated transcription factors (TFs), and ATAC narrowPeak BED file to search for TF motifs within super-enhancer regions. Remaining parameters were left as default. Clique fractions were calculated as the number of cliques a TF participates in divided by the number to total cliques.

## ACKNOWLEDGMENTS

We thank staff from the Animal Facility, Genotyping Core, Flow Cytometry Facility, Molecular Genomics Core, Victorian Center for Functional Genomics (VCFG), Centre for Advanced Histology and Microscopy (CAHM) and Bioinformatics Consulting Core (BCC) of the Peter MacCallum Cancer Centre, and members of the Johnstone laboratory for useful discussions. We acknowledge Nicole Messina and Dan Andrews for providing reagents and advice and support from the Peter MacCallum Cancer Centre Foundation and Australian Cancer Research Foundation.

Research reported in this publication was supported by Cure Cancer Australia under award number 1051444. S.J.V. was supported by a Rubicon Fellowship from the Netherlands Organization for Scientific Research (NWO, 019.161LW.017), an NHMRC EL1 Fellowship (GNT1178339) and a Peter MacCallum Cancer Foundation Grant; S.J.H. was supported by a Postdoctoral Fellowship from the Cancer Council of Victoria (CCV); L.M. and SCIL were supported by The Lorenzo and Pamela Galli Research Fund Trust; S.S. by the German Research Foundation (DFG – 270650915/GRK2158 and SCHE692/6-1) and by the Manchot Graduate School ‘Molecules of Infection III’; D.D.C. is funded by Canadian Institute of Health Research (201512MSH360794-228629, FDN 148430, PJT 165986) and by the Canada Research Chairs; B. T. K. by a Project Grant from NHMRC (Grant No. 1113577) and a NHMRC Principal Research Fellowship (No. 1063008); R.J.W by a Project Grant from Cancer Council Victoria, a Project Grant and Program Grant (Grant 454569) from the NHMRC, a NHMRC Senior Principal Research Fellowship and a Grant from The Kids’ Cancer Project (to R.W.J. and S.J.V).

---

**Supplementary Fig. 1.**
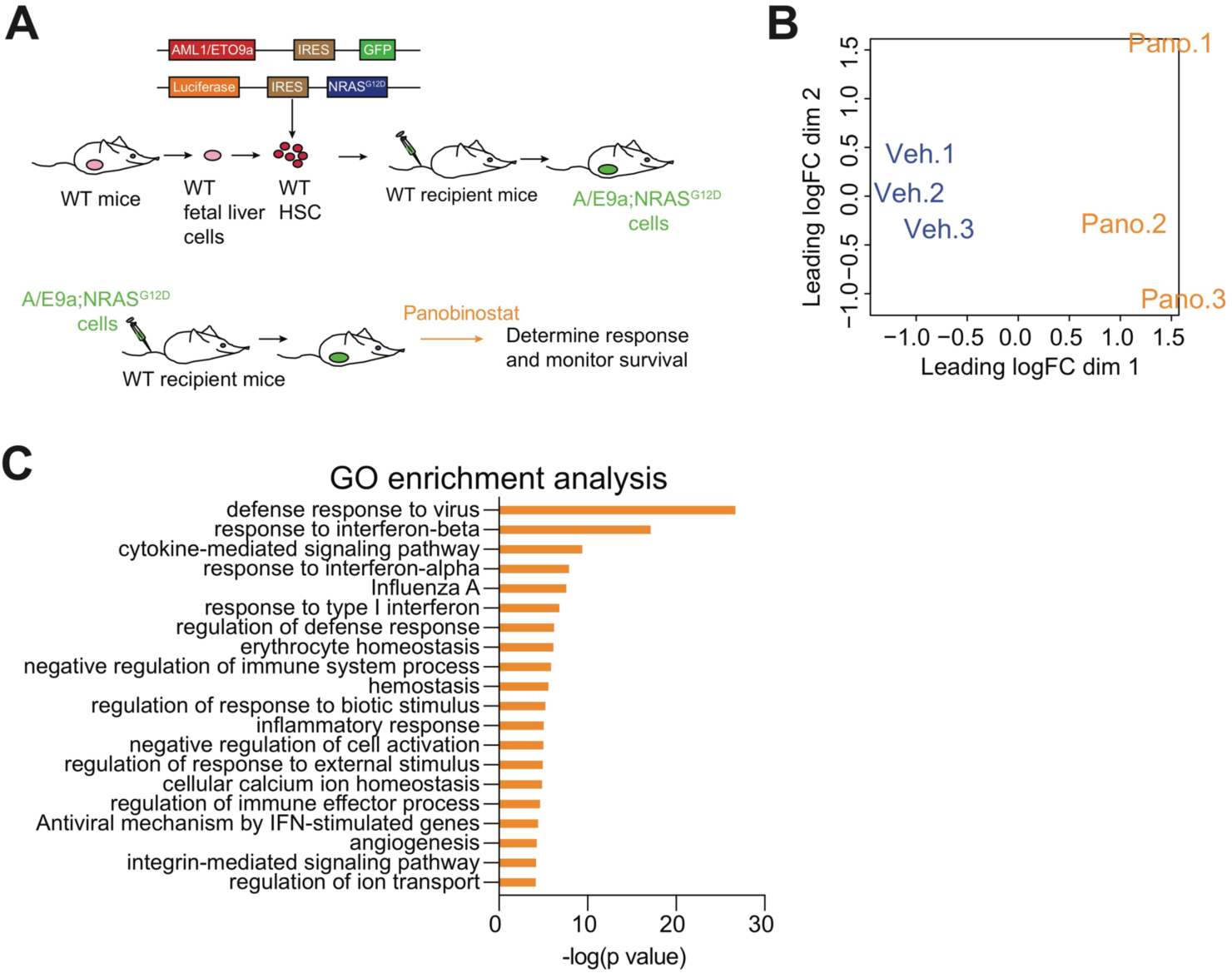
(**A**) Schematic representation of the A/E9a;NRAS^G12D^ leukemia model. (**B**) Principal component analysis of RNA-seq data of A/E9a;NRAS^G12D^ leukemia cells from panobinostat-treated and vehicle-treated mice. (**C**) Pathway enrichment analysis of upregulated genes between panobinostat-treated and vehicle-treated A/E9a;NRAS^G12D^ leukemia cells.

**Supplementary Fig. 2.**
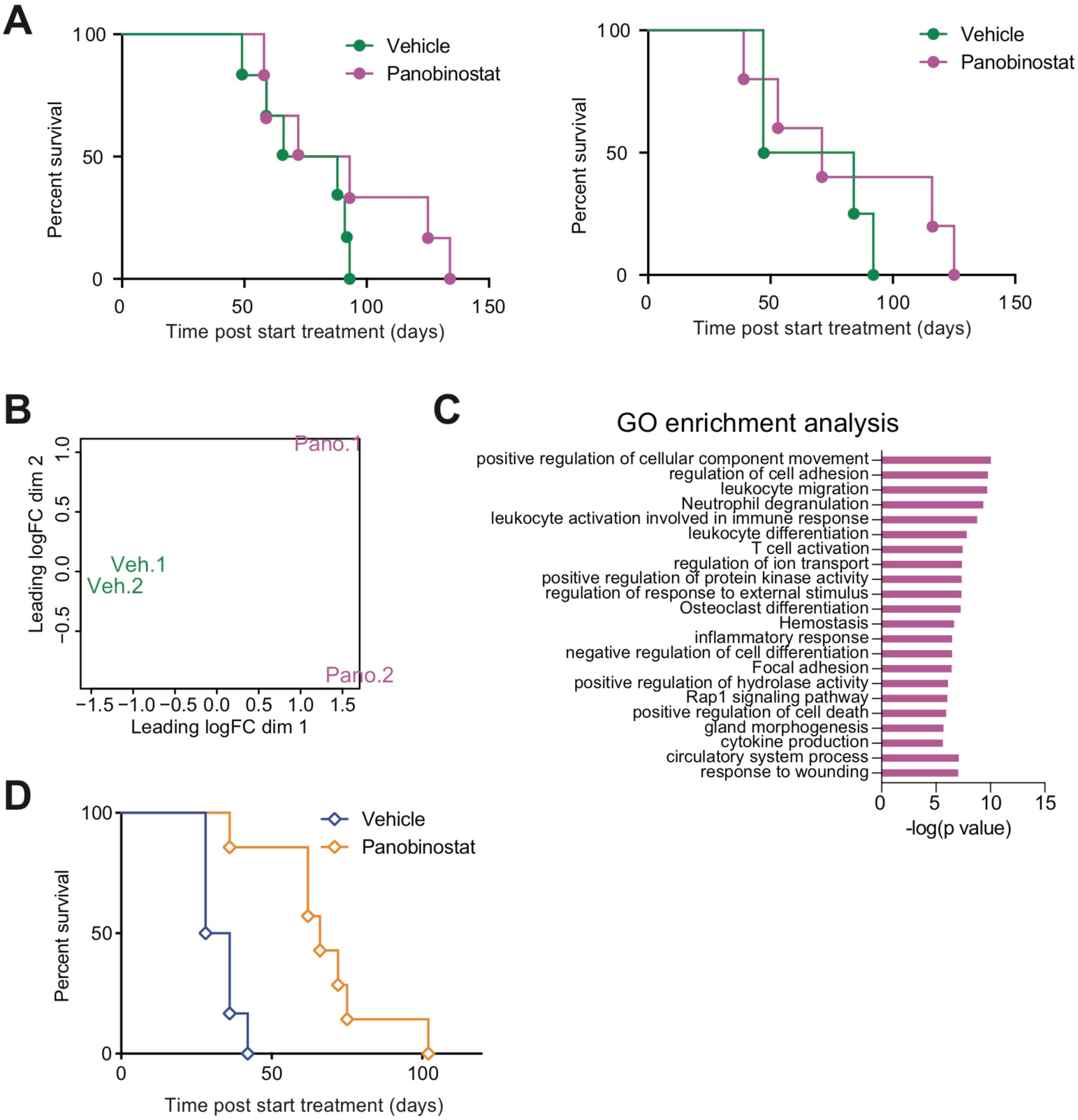
(**A**) Kaplan-Meier survival curves of mice bearing A/E9a;NRAS^G12D^;*Ifnar1*^-/-^-driven leukemias (left: tumor clone 1.3; right: tumor clone 3.3) treated with either vehicle or panobinostat (n=4-6 mice per treatment group). (**B**) Principal component analysis of RNA-seq data of A/E9a;NRAS^G12D^;*Ifnar1*^-/-^ leukemia cells from panobinostat-treated and vehicle-treated mice. (**C**) Pathway enrichment analysis of upregulated genes between panobinostat-treated and vehicle-treated A/E9a;NRAS^G12D^;*Ifnar1*^-/-^ leukemia cells. (**D**) Kaplan-Meier survival curves of *Ifnar1*^-/-^ mice bearing WT A/E9a;NRAS^G12D^*-* driven leukemias and treated with either vehicle or panobinostat (n=6-7 mice per treatment group)

**Supplementary Fig. 3.**
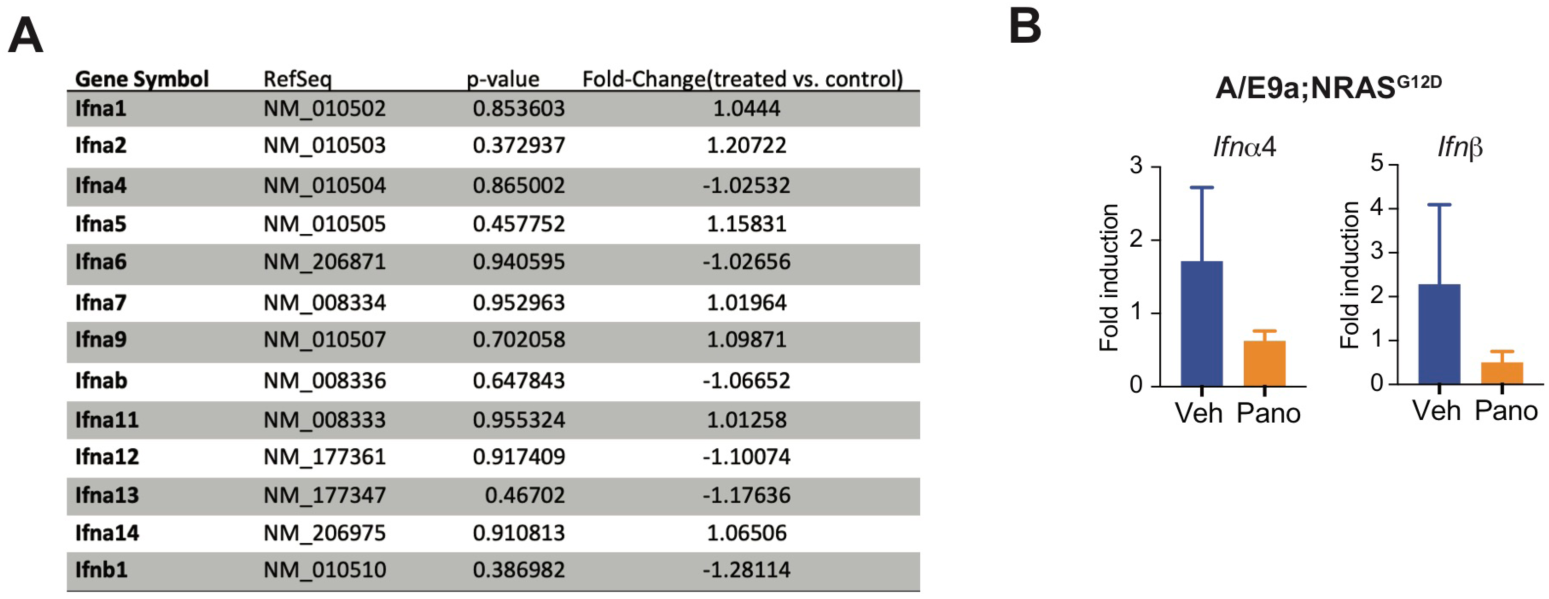
(**A**) Microarray values for *Ifn*α isoforms and *Ifn*β mRNA in tumor cells isolated from the spleen of mice bearing A/E9a;NRAS^G12D^-driven leukemias treated with either vehicle (control) or panobinostat following 5 days of therapy (**B**) qRT-PCR of *Ifn*α*4* and *Ifn*β transcripts in tumor cells isolated from the spleen of mice bearing A/E9a;NRAS^G12D^-driven leukemias treated with either vehicle or panobinostat following 5 days of therapy.

**Supplementary Fig. 4.**
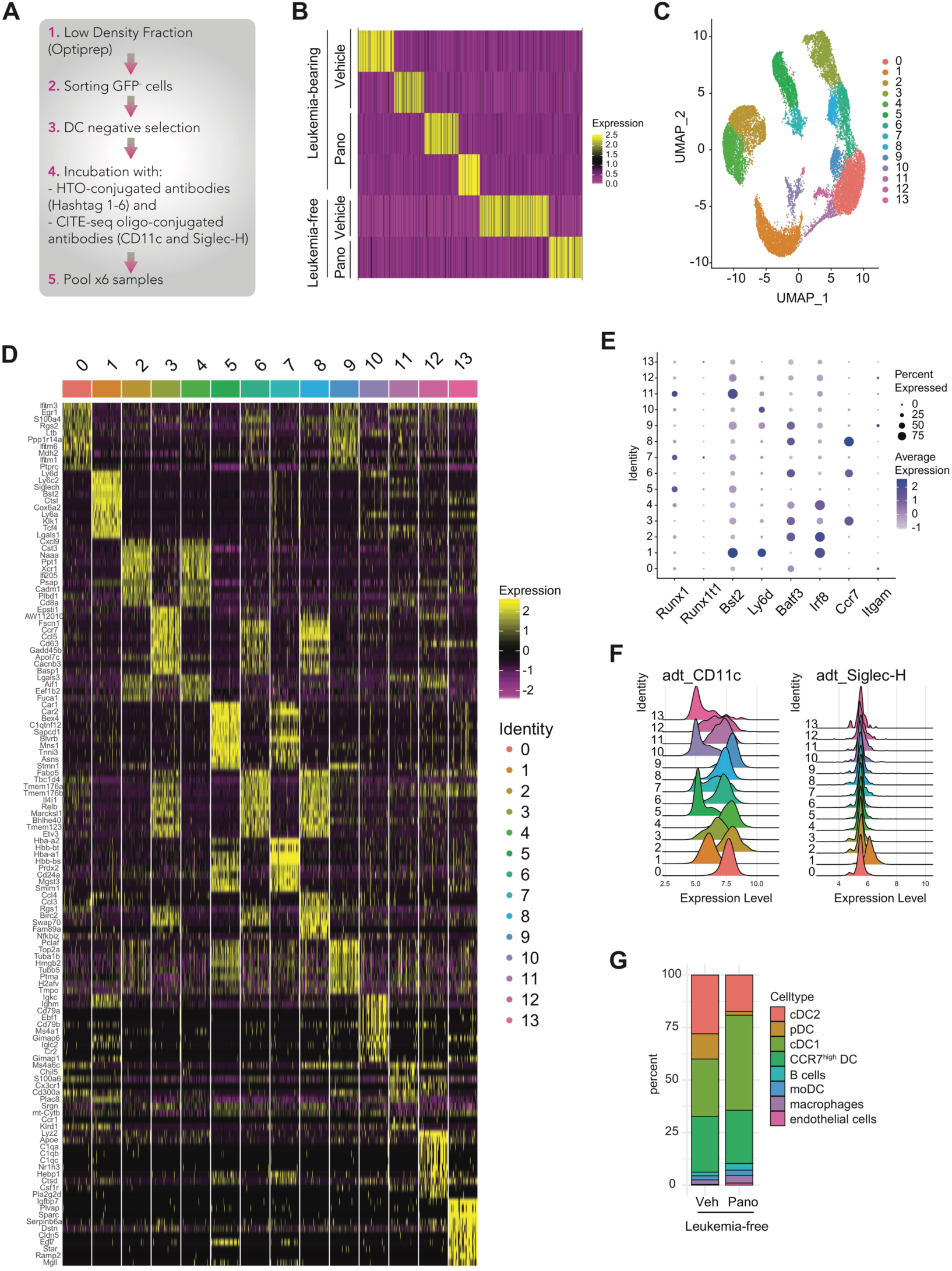
(**A**) Multi-step preparation of splenic DC suspensions for sc-RNAseq. (**B**) Heatmap showing proportion of HTO for each sample: A/E9a;NRAS^G12D^-vehicle (x2), A/E9a;NRAS^G12D^-panobinostat (x2), WT-vehicle (x1), WT-panobinostat (x1). (**C**) Total number of cell clusters based on RNA expression. (**D**) Gene expression of individual clusters. (**E**) Expression of selected genes in individual clusters. (**F**) Expression of CD11c and Siglec-H proteins in individual clusters. (**G**) Relative frequency of individual clusters, excluding tumor/myeloid cells.

**Supplementary Fig. 5.**
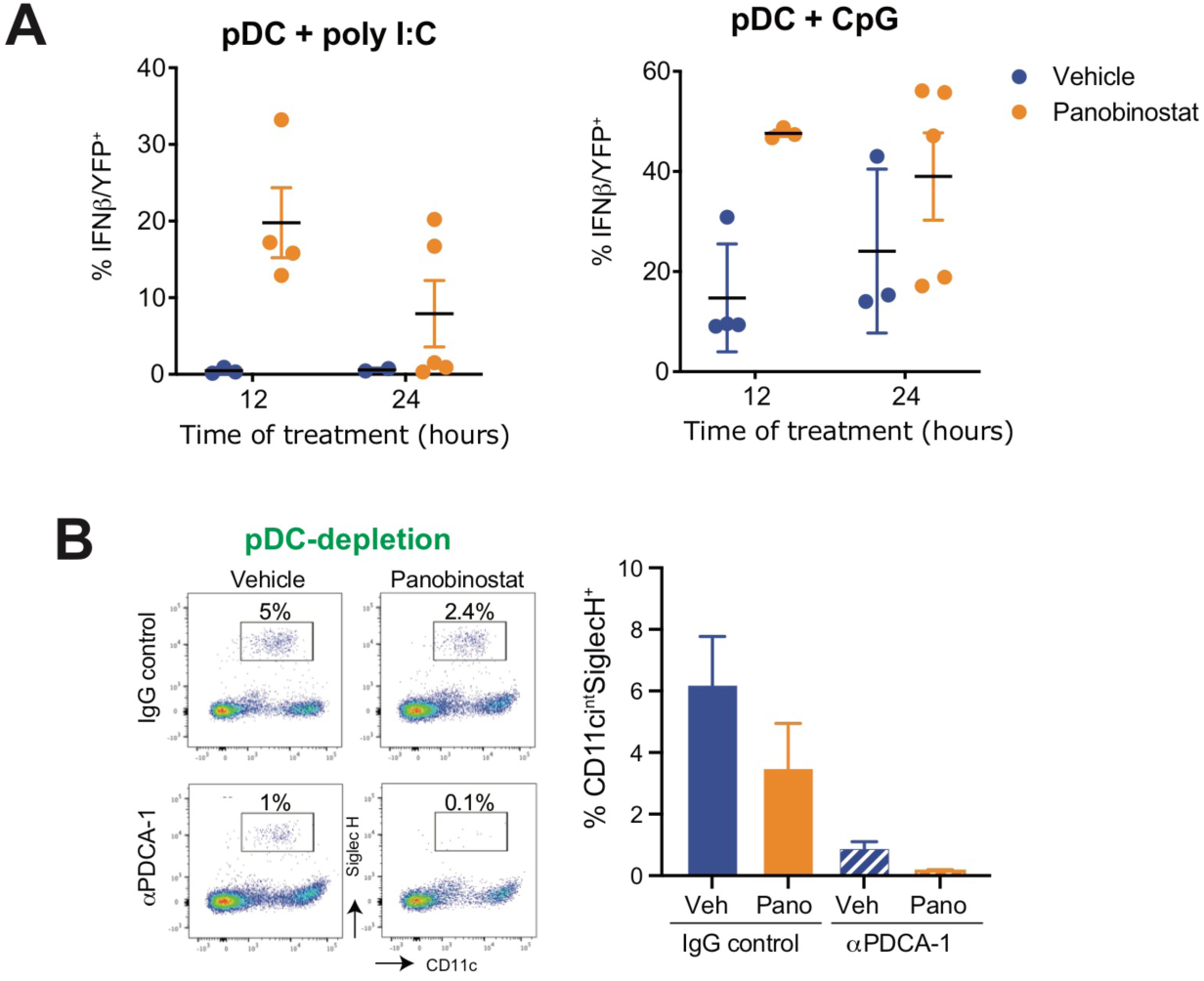
(**A**) Percentage of IFNβ/YFP^+^ pDCs gated from *ex vivo* bone marrow cells stimulated with poly I:C (left) or CpG (right) for the indicated time in combination with panobinostat (or vehicle as control). (**B**) Flow cytometry plot (left) and percentage (right) of pDC (CD11c^int^SiglecH^+^) in mice bearing A/E9a;NRAS^G12D^-driven leukemias receiving IgG control or anti-PDCA-1 and treated with either vehicle or panobinostat following 3 days of therapy.

**Supplementary Fig. 6.**
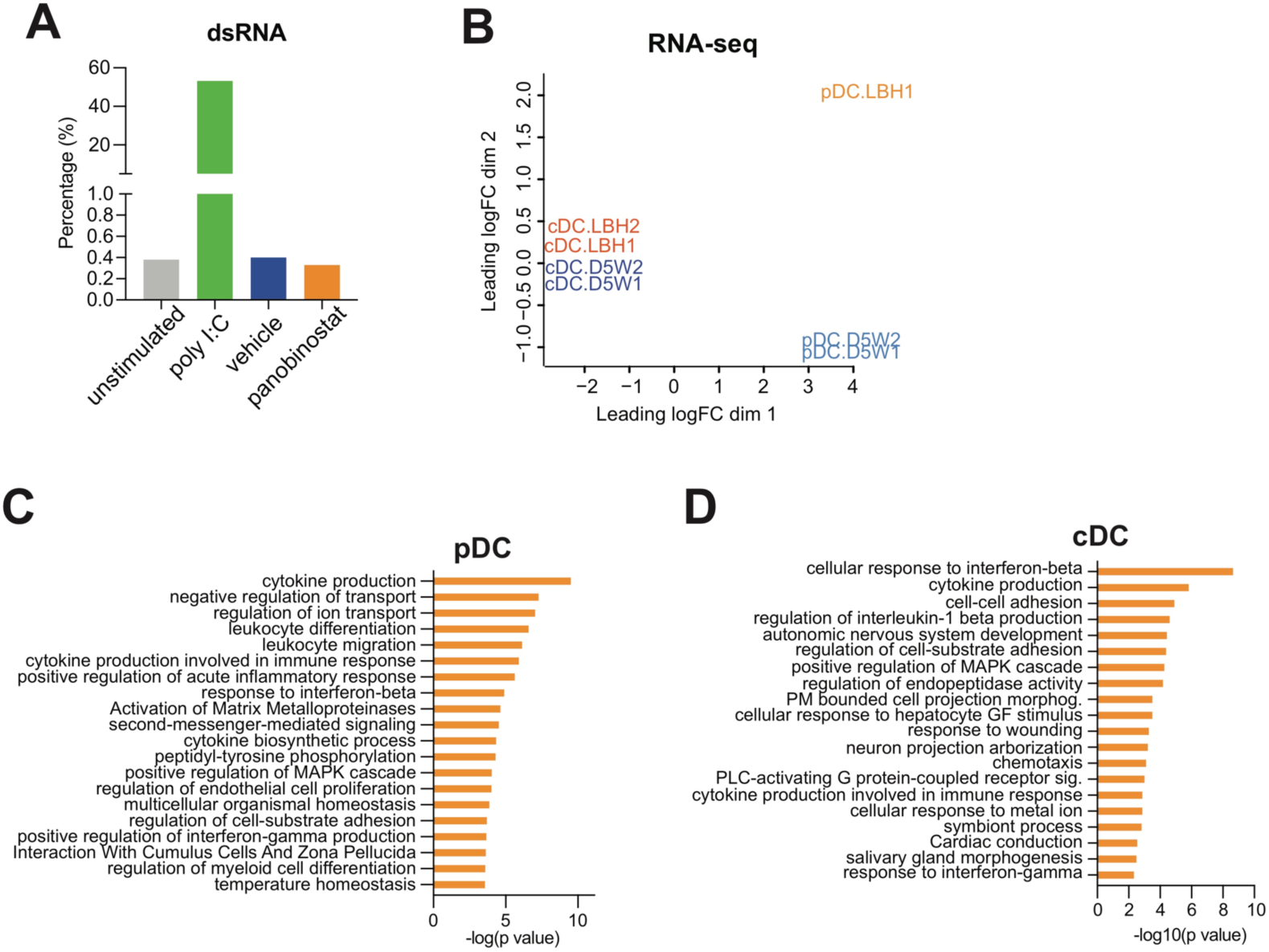
(**A**) Detection of dsRNA in Flt3L-derived pDC treated with vehicle or 16nM panobinostat for 6h. As a positive control, cells were stimulated with transfected poly(I:C), a synthetic analogue of dsRNA which is sensed by RIG-I/MDA-5. (**B**) Principal component analysis of RNA-seq of pDC and cDC isolated from panobinostat-treated and vehicle-treated A/E9a;NRAS^G12D^ leukemia-bearing mice. (**C-D**) Pathway enrichment analysis of upregulated genes (pano vs vehicle) in pDC (C) and cDC (D) isolated from A/E9a;NRAS^G12D^ leukemia-bearing mice treated with vehicle or panobinostat for 2 days.

**Supplementary Fig. 7.**
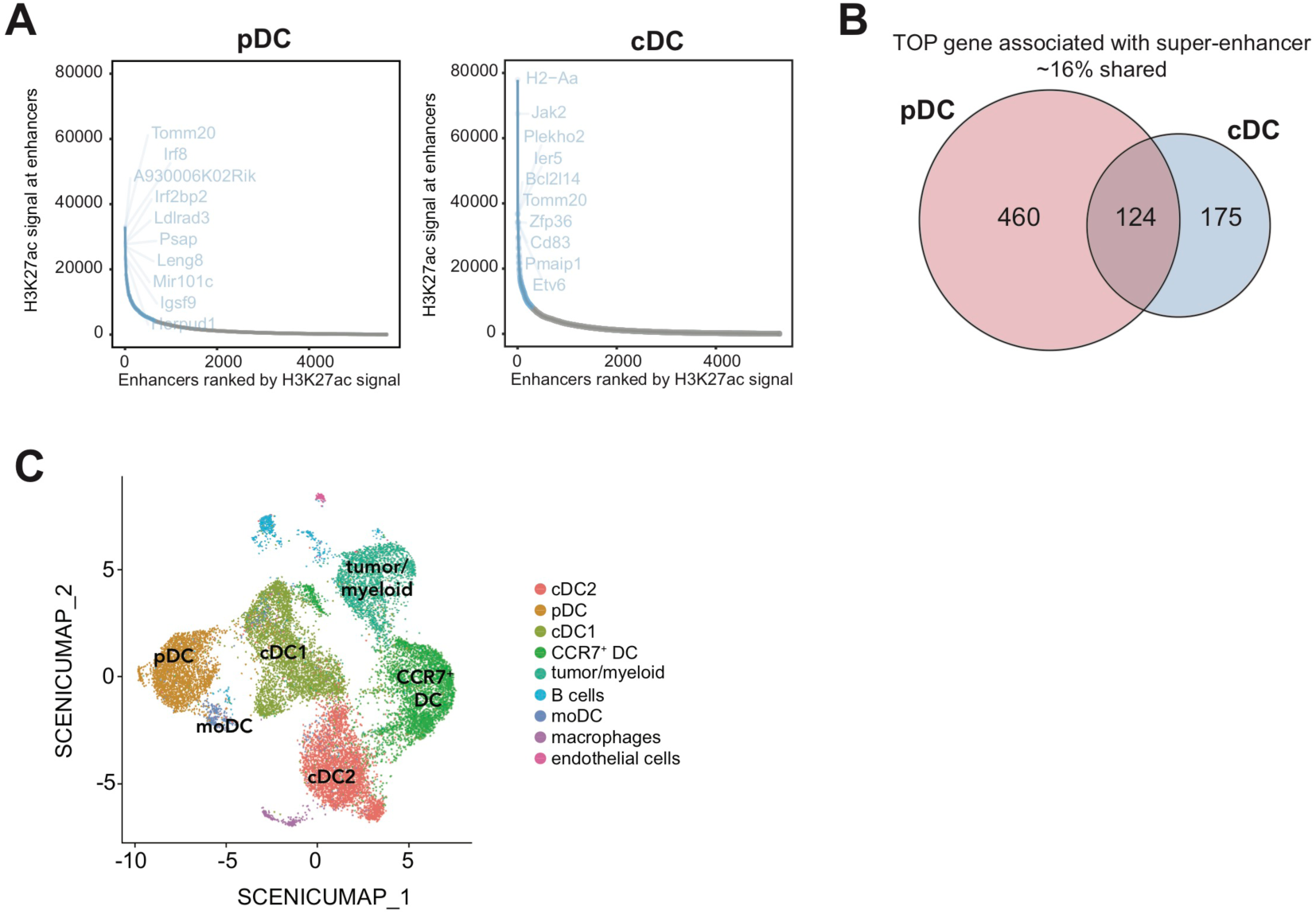
**(A)** Ranked enhancer plots defined across H3K27ac landscapes of pDC and cDC. Genes associated with SE are highlighted. **(B)** Venn diagram showing TOP genes associated with SE in pDC and cDC. (**C**) UMAP visualization of SCENIC-defined cell clusters.

